# Late maturation of semantic control promotes conceptual development

**DOI:** 10.1101/2022.03.31.486559

**Authors:** Rebecca L. Jackson, Matthew A. Lambon Ralph, Timothy T. Rogers

**Affiliations:** Department of Psychology & York Biomedical Research Institute, University of York, Heslington, York, YO10 5DD; MRC Cognition & Brain Sciences Unit, University of Cambridge, 15 Chaucer Road, Cambridge, CB2 7EF; 524 WJ Brogden Hall, Department of Psychology, University of Wisconsin-Madison, Madison, Wisconsin, 53706

**Keywords:** executive control, semantic cognition, semantic control, neural networks, development, learning

## Abstract

Control processes underpinned by the prefrontal cortex are critical for generating task-appropriate behaviour across cognitive domains, yet this region develops extremely late. Traditionally, this developmental pattern is considered negative but necessary. However, an alternative (yet perhaps complementary) view suggests that a developmental period without control could support learning, particularly in the semantic domain. Here, we exploit a recent computational model to test formally whether late development of the context-sensitive use of conceptual knowledge, or ‘semantic control’, would promote concept acquisition. Simulations show that late maturation of semantic control and anatomical connectivity conspire to promote conceptual learning. Delayed control speeds conceptual learning without compromising conceptual representations, particularly when semantic control connects to intermediate layers. To assess whether semantic control also develops late in human children, we conducted a meta-analysis of the classic triadic matching task where participants decide which of two options best matches a third. Matching can be based on taxonomic or thematic relations. When these conflict, participants must exert semantic control to determine which relation is task appropriate. Context-sensitivity develops later than conceptual knowledge with large increases between 3 and 6 years. Thus, the protracted PFC development leads to a delay in acquiring semantic control processes, benefiting conceptual learning.

## Introduction

Human thought is remarkably flexible. More than other mammals, humans can control the mental representations and behaviours that arise from moment to moment, avoiding prepotent responses and reconfiguring well-learned patterns to better suit immediate and long-term goals. These abilities depend upon the gradual maturation of the prefrontal cortex (PFC; Duncan, 2010; Fuster, 2015; Miller & Cohen, 2001; Smaers, Gómez-Robles, Parks, & Sherwood, 2017; Wood & Grafman, 2003), a brain structure that develops more slowly than other cortical regions within the human brain and more slowly in humans than the homologous structures of other mammals (Chugani & Phelps, 1986; Elston, Oga, & Fujita, 2009; Huttenlocher & Dabholkar, 1997). Slow developmental change is observed in structural (synaptogenesis, synaptic elimination, dendritic arborisation and myelination) and functional (regional metabolism) measures, with the PFC not fully maturing until past the age of twenty years (Casey, Tottenham, Liston, & Durston, 2005; A. Diamond, 2002; Gogtay et al., 2004; Mrzljak, Uylings, van Eden, & Judas, 1990; Sowell, Trauner, Gamst, & Jernigan, 2002; Veroude, Jolles, Croiset, & Krabbendam, 2013). As a result, behavioural tasks of domain-general executive function (speed of processing, working memory, inhibition and task switching) mature slowly across childhood and adolescence with children undergoing a long developmental period with comparatively little executive control (Davidson, Amso, Anderson, & Diamond, 2006; A. Diamond & Doar, 1989).

The slow onset of domain-general executive control is often viewed as an unfortunate but necessary consequence of the PFC’s complexity (e.g., Church, Bunge, Petersen, & Schlaggar, 2017; A. Diamond, 2002; Zelazo, Carlson, & Kesek, 2008). Given constraints on the size of the head at birth, the requirement of growing out and pruning back connections from PFC to the rest of the brain, and other complex biological constraints, it may be that control processes simply require a very long time to mature (Bunge, Dudukovic, Thomason, Chandan, & Gabrieli, 2002; Church et al., 2017; Crone, Donohue, Honomichl, Wendelken, & Bunge, 2006; Luna, Garver, Urban, Lazar, & Sweeney, 2004; Wilk & Morton, 2012). We consider an alternative, though perhaps complementary view: that the slow maturation of the PFC, and therefore associated control processes, has positive functional consequences for learning, which offset the costs incurred by the late development of flexible and adaptive behaviour. Building on prior proposals (Chrysikou, Novick, Trueswell, & Thompson-Schill, 2011; Newport, 1990; Thompson-Schill, Ramscar, & Chrysikou, 2009), we specifically consider that later development of control may confer a functional advantage in the development of conceptual knowledge (or ‘semantics’). As such we focus on the development of ‘semantic control’, the context-sensitive use of conceptual knowledge to perform a current task or goal (Jackson, 2021; Jefferies, 2013; Lambon Ralph, Jefferies, Patterson, & Rogers, 2017; Reilly et al., 2024). Unlike domain-general executive control, the developmental trajectory of semantic control is unknown, allowing us to first consider whether it would be beneficial for semantic control to develop later than semantic representation and then subsequently, to consider whether it does develop late in human development.

While many domains require control processes to shape behaviour, semantic control relies on at least partially distinct systems from executive control more generally. Functional brain imaging and patient studies suggest that the ventrolateral prefrontal and posterior temporal substrates crucial for semantic control are specialised for the control of meaningful content, and not domain-general executive control (Chiou, Jefferies, Duncan, Humphreys, & Lambon Ralph, 2023; Gao et al., 2021; Hodgson, Lambon Ralph, & Jackson, 2021; Hodgson, Lambon Ralph, & Jackson, 2024; Humphreys & Lambon Ralph, 2017; Jackson, 2021; Jefferies & Lambon Ralph, 2006). Thus, the regions supporting semantic control are at least partially dissociable from those supporting the executive control of other domains, with differential reliance on ventrolateral versus dorsolateral prefrontal areas, as well as posterior temporal versus parietal cortices. Such dissociations may partially reflect the differences in how control operates in semantic retrieval versus other executive tasks. Semantic control makes use of contextual information to generate accurate and task-appropriate semantic inferences—for instance, inferring the weight of a piano rather than its function in the context of moving house, but function rather than weight in the context of a recital. In both cases, the information retrieved is true of the piano, and the control system shapes which inference is to guide subsequent behaviours in a context-appropriate manner. Neither fact about pianos is arbitrary, unfamiliar, or difficult to retrieve; they are simply relevant in different situations.

In contrast, executive control tasks often employ measures requiring the inhibition of some strong behavioural impulse, or a tendency to behave in accordance with a novel, arbitrary rule (e.g., Kochanska, Murray, & Harlan, 2000)—a kind of behaviour known to develop late (e.g., Zelazo et al., 2008). For instance, several tasks require the child to forego an immediate reward for a period of time (e.g., Delay of Gratification, Forbidden Toy, Gift Delay, Pinball, Snack Delay), to abandon impulsive behaviours in favour of those that accord with social conventions (e.g., Disappointing Gift; Whisper task; turn-taking in the Tower task), to inhibit previously successful behaviours (e.g. multilocation search tasks, dimensional-change card sort) or responses to perceptually salient stimuli (e.g., shape Stroop), or to generate temporal sequences of behaviours conforming to an arbitrary rule (e.g., backward digit span, count-and-label, motor sequencing). While some executive tasks do include selection and inhibition processes that seem to parallel those of semantic control (such as the Cattell Culture Fair Task; Cattell, 1971; Woolgar, Bor, & Duncan, 2013), nevertheless such tasks recruit different brain networks for meaningful words versus meaningless shape stimuli even when task processes and demands are matched, with greater ventrolateral PFC and posterior temporal cortex involvement for meaningful stimuli (Hodgson et al., 2024).

Thus, delineating the developmental trajectory of semantic control specifically requires tasks in which control is applied to the context-appropriate retrieval and manipulation of semantic knowledge. Unlike domain-general executive control, the development of semantic control has not been extensively studied. Because semantic control also relies on the PFC, it seems plausible that it should follow a similar developmental trajectory to general executive function. Yet the PFC itself is neither anatomically nor functionally homogeneous, with different subregions maturing along different timelines (Gogtay et al., 2004; Sowell et al., 2002). Because semantic control relies upon a distinct ventrolateral PFC region, it is equally possible that its developmental trajectory differs from that of domain-general executive control.

In the rest of this paper, we explain why conceptual knowledge acquisition in particular, might benefit from such a period with limited control, assess the hypothesis formally via simulations with a computational model of controlled semantic cognition, and then test for evidence of such a delay in semantic control by reconsidering classic findings in studies of conceptual development in light of this perspective and its simulated advantages.

### Central idea and motivation from prior work

Our proposal is motivated by the seemingly antagonistic relationship lying at the heart of controlled semantic cognition (Jackson, Rogers, & Lambon Ralph, 2021; Lambon Ralph et al., 2017). On one hand, the human semantic system must accumulate information about the world across many disparate learning episodes, each providing only a partial glimpse into the environment’s structure. Consider the widespread view that human concepts arise from learning bundles of correlated attributes in the environment (Keil, 1979; Murphy & Medin, 1985; Rogers & McClelland, 2004; Rosch, Mervis, Gray, Johnson, & Boyes-Braem, 1976). For instance, the concept *bird* might arise because there are sets of attributes that all tend to occur together in various instances involving birds: feathers, wings, beaks, hollow bones, the ability to fly, the ability to lay eggs, the name “bird,” and so on. While these properties may all be true of the same items, such covariation is not transparently observable in everyday experience: a flying bird is not simultaneously laying an egg; the bird skeleton observed in science class does not possess feathers and does not fly; the word “bird” appearing in a novel typically does not appear together with an image of the bird, and so on. To learn that these various properties all inhere together within exemplars of the concept *bird*, the semantic system must be able to track sameness of kind across these different scenarios—that is, it must “know” that the bird observed flying overhead is the same kind of thing as the skeleton studied in science class, which in turn is the same kind of thing as the item described in a novel. Thus, the semantic system must *abstract transtemporally* across different situational contexts, each providing only a partial glimpse into an item’s full complement of attributes, to form conceptual representations that fully express attribute covariances as they exist in the world and allow generalisation to new exemplars (Jackson et al., 2021; Lambon Ralph et al., 2017; Rogers & McClelland, 2004). On the other hand, to support everyday behaviour the semantic system must also produce internal representations and behaviours that are *context-sensitive,* so that information that comes to mind in the moment is suited to the goals of the immediate behavioural task. The features required to engage with a piano in different contexts, such as moving it and playing it, differ substantially (Saffran, 2000). In order to provide a context-appropriate output, only relevant features should be selected and allowed to bear on the output decision whilst other, often more dominant, features must be suppressed (Badre, Poldrack, Pare-Blagoev, Insler, & Wagner, 2005; Jefferies, 2013; Thompson-Schill, D’Esposito, Aguirre, & Farah, 1997; Wagner, Pare-Blagoev, Clark, & Poldrack, 2001). Thus, the semantic system must concurrently abstract across contexts to acquire representations that express conceptual structure (*semantic representation*), yet also shape representations and behaviours to suit the immediate context (*semantic control*).

We previously described a neural network model that simulates both semantic control and representation (Jackson et al., 2021). Because this model learns over time via simulated experiences with various objects across various tasks, it provides a means of investigating the developmental trajectories of semantic control and representations under different hypotheses. Semantic control signals are thought to promote the activation of task-relevant properties while suppressing task-irrelevant properties true of a given item (Cohen, Dunbar, & McClelland, 1990; Thompson-Schill et al., 1997; Waksom, Kumaran, Gordon, Rissman, & Wagner, 2014). For instance, when naming an item such as a toy train, semantic control ensures that only a context-appropriate name is produced (e.g., “train” and not “vehicle,” “toy,” or “choo-choo”), and moreover that other associated information, such as action-plans (e.g., playing with the toy) or other irrelevant facts about the object (“It goes on tracks”) do not drive behaviour. Thus, the model of controlled semantic cognition must learn concepts by integrating their constituent features across sensory modalities, episodes and contexts, reflecting semantic representation processes, but it must also produce context-sensitive responses based on current task goals: features presented as input in one of three informational modalities must generate output across some but not all modalities based on a task context representation originating in PFC. This semantic control demand reflects the selection of task-appropriate and/or inhibition of task-irrelevant conceptual features necessary for context-sensitive behaviours. As it may be counterproductive to become aware of aspects of a concept that would be irrelevant, or even actively unhelpful in making a response, such guided activation is considered a necessary precursor to response selection (Botnivick & Cohen, 2014).

Prior work with this framework compared multiple different architectures, reflecting different hypotheses about the structure of the semantic network, in their ability to abstract conceptual structure across modalities and contexts even while producing context-sensitive outputs. We found that these opposing functional demands are best accomplished within an architecture like that shown in Figure 1, where representations across different modalities (e.g., vision, audition, language, action) mutually interact via a single deep, multimodal “hub” (Jackson et al., 2021). In this architecture, the “hub” units are always involved in propagating information between different “spokes” of the system, and so can learn the high-order covariance structure of the environment across different modalities and contexts—even though, for each training episode, the control system allows only a subset of an item’s properties to become active.

**Figure 1.**
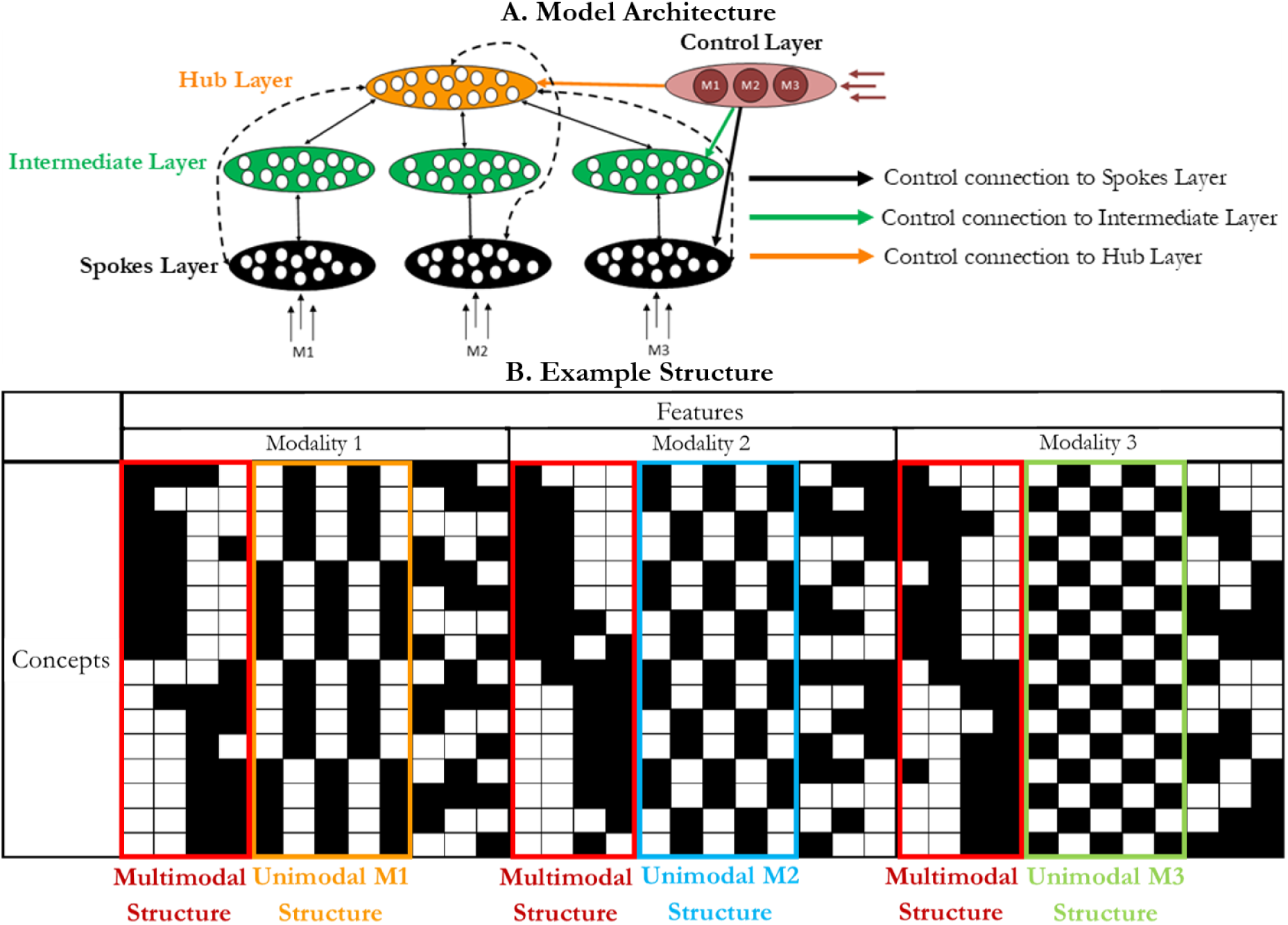
A. The model architecture. Sensory input is received, and output generated, in three modality-specific sensory regions (black ovals) constituting the Spokes Layer. This layer is connected to an Intermediate Layer (green ovals) which in turn is connected to the Hub Layer (orange oval). A small number of sparse connections also directly connect the Spokes Layer to the Hub Layer. A context signal is received as input to three units forming a Control Layer (red oval). This context signal specifies the modality in which the sensory input is received and the modality in which the output should be generated. This Control Layer may be connected to all layers (full connectivity), or to the Spokes Layer (black arrow), Intermediate Layer (green arrow) or Hub Layer (orange arrow) only. B. The model environment. The 16 concepts are shown. Each concept is a row, and each feature is a column. If the feature is present the box is black, if absent the box is white. When control requirements are present, each concept can be presented in one of three modalities, and output required in one of three modalities, resulting in 144 different trials. Each modality has a surface similarity structure (orange, yellow or green boxes), which is orthogonal to a multimodal structure (red boxes) learnt through conceptual abstraction across modalities.

Furthermore, semantic representation and control were best supported by an architecture where control either operates directly on task-relevant modality-specific regions or on immediately adjacent areas (corresponding to the presence of the black or green but not orange connections in Figure 1; Jackson et al., 2021). That is, when the multimodal hub is insulated from the direct effects of control, the representations that emerge there are relatively invariant to context, and thus come to express the conceptual structure inherent across different learning episodes regardless of the current contextual demands. Thus, the antagonistic processes of control and representation can co-exist in a semantic system with an architecture that promotes their relative functional separation.

This “reverse engineered” model architecture also conformed well to the organisation of the cortical semantic system, and successfully accounted for key neuroimaging and neuropsychological data. For instance, the systems of semantic control versus representation demonstrate relative anatomical segregation within the brain, with PFC and lateral posterior temporal cortex supporting semantic control processes, and the anterior temporal lobes supporting multimodal semantic representation (Jackson, 2021; Jefferies, 2013; Lambon Ralph et al., 2017; Patterson, Nestor, & Rogers, 2007; Rogers et al., 2004). This organisational principle also underlies the distinct behavioural profiles observed in *semantic dementia*—a gradual erosion of multimodal conceptual representations accompanying a progressive degradation of the anterior temporal lobe (Acosta-Cabronero et al., 2011; Patterson et al., 2007; Warrington, 1975)—and *semantic aphasia*, a loss of semantic control or the controlled, context-appropriate use of concepts following cerebrovascular accident to frontal or posterior temporal cortices (Jefferies, 2013; Jefferies & Lambon Ralph, 2006), a neuropsychological dissociation well captured by the reverse-engineered model.

### Importance of development

This paper considers whether late maturation of semantic control might likewise promote better acquisition of conceptual structure, and whether such effects obviate or complement prior conclusions about the optimal anatomical connectivity between semantic control and representation regions. The model can be required to produce context-sensitive responses from the start of learning (as in prior work), or after a time period in which the model does not tailor its behaviour to the context, allowing us to evaluate whether and how the length of this delay influences the system’s ability to both acquire conceptual structure and generate, at maturity, context-sensitive outputs.

To understand why this might be, consider that controlled responding requires the suppression of task-irrelevant properties and/or potentiation of task-relevant properties. Since conceptual structure is latent in the distribution of properties across tasks, it may be difficult to learn when only task-relevant properties are permitted to activate from the outset of development. Moreover, conceptual acquisition may be generally slowed in this situation, since the learner must inhibit all but a small subset of an item’s properties in any given episode. Thus, a developmental period with limited control may promote conceptual learning because the absence of control allows all properties associated with a stimulus, including those not relevant to the current task and not directly observed, to become activated simultaneously.

Moreover, these effects could differ based on the overall architecture of the system. If delayed onset of control promotes conceptual abstraction, it may be less important for the control system to be anatomically insulated from the multimodal semantic hub, obviating conclusions from prior work that the control and hub regions should not directly connect.

Alternatively, it may be that the architectural constraints identified in this prior work are sufficient to ensure good learning of conceptual structure regardless of the timing with which the control system develops, or that the developmental timing and architectural constraints conspire to best promote conceptual abstraction. To adjudicate these possibilities, the first part of this paper reports simulations with the reverse-engineered model of controlled semantic cognition (Jackson et al., 2021), manipulating both the rate at which the control system “matures” and the connectivity between the control and semantic representation networks, then measuring the effect of these factors on conceptual abstraction and training time. The simulations provide clear conclusions about both the development of semantic control and the connectivity of semantic control and representation regions.

The second part of the paper then considers predictions of the simulations about the time-course of semantic control over children’s development. The development of both executive function (e.g., Davidson et al., 2006; A. Diamond & Doar, 1989) and of semantic representation (e.g., Gershkoff-Stowe & Rakison, 2005; Keil, 1979; Mandler & McDonough, 1993; Zelazo et al., 2008) have been extensively studied. However, as noted above, comparatively little work has touched on questions about semantic control specifically (Blaye, Bernard-Peyron, Paour, & Bonthoux, 2006; Blaye & Jacques, 2009; Henderson, Clarke, & Snowling, 2011; Snyder & Munakata, 2010, 2013) therefore the relative time-course under which semantic information becomes available to the developing child versus when this information can be used in a controlled fashion remains unclear.

Part 2 briefly reviews this literature, then evaluates the model predictions through a new meta-analysis of a classic task designed to test development of semantic knowledge: the triadic match-to-sample task in which children must decide which of two items is most similar to a third. While this task has not traditionally been viewed as assessing semantic control, many studies have focused on manipulating both the nature of the information required for a successful match (taxonomic or thematic; Bauer & Mandler, 1989; Berger & Aguerra, 2010; Blanchet, Dunham, & Dunham, 2001; Greenfield & Scott, 1986; Smiley & Brown, 1979; Waxman & Namy, 1997) and the influence of instructional cues on behaviour (e.g., Estes, Golonka, & Jones, 2011; Greenfield & Scott, 1986; Waxman & Namy, 1997). As an instructional cue is a form of task context, sensitivity to the cue—that is, the degree to which children can make different choices for the same stimulus depending upon the task instruction—indexes the ability to use semantic control to shape semantic decisions. Our meta-analysis tracks changes in such sensitivity over development. Revisiting this extensive literature through the contemporary lens of controlled semantic cognition then provides a powerful way to estimate the interplay of semantic control and representation processes across development.

#### Part 1: Simulations

We conducted simulations with the recent model of controlled semantic cognition from Jackson et al., (2021; see Figure 1), a deep variant of the hub-and-spokes framework in which the propagation of activation through the semantic representation network is constrained by inputs from a prefrontal “control” system. Items in the environment are represented as distributed patterns across three modality-specific input/output channels (e.g., vision, words, and action) and the model always receives the features of a concept in one sensory channel as input. In the full “mature” model control requirements are present, meaning the model receives a task context signal dictating a required output modality and learns to produce output patterns in the channels relevant to the current task context only. Thus, the trained model must activate only properties that are both true of the item and relevant to the current task context. However, the model can be trained initially without this requirement. Here we introduced an initial *control-free* period where no task context signal was provided and where the model was trained to output all features relevant to a concept regardless without a specific task context (as in prior models simulating semantic representation only). This is analogous to rewarding a child for item-relevant responses regardless of their task relevance, such as rewarding either the answer “choo choo” or “train” (but not, for example, “moo” or “cow”) when tasked with naming a train. After a variable period of time, controlled training items were introduced with gradually increasing frequency allowing the model to develop into the mature system, meeting the central challenges of controlled semantic cognition by responding with only the features both true of the item and relevant to the task.

To understand how control influences the development of semantic representations, we manipulated the timing and speed with which controlled semantic behaviour arises. Specifically, we varied (a) the *maturational delay*, that is, the duration of the training period without semantic control, ranging from 0 to 5000 epochs in 1000 epoch increments, (b) the *transition function* describing how the frequency of controlled learning trials increased over time (linear or sigmoidal), and (c) the size of the *transition window* over which controlled trials ramped up from 0-100% of training trials (500, 1000, or 2000 epochs). We also assessed models which maintained no control or full control throughout for comparison only.

To understand how the connectivity and maturation of semantic control impact simulated conceptual development, we considered two metrics. The first is *training time,* measured as the number of learning epochs required before the model generates the correct task-specific outputs for all *controlled* patterns in the environment. The second is the *conceptual abstraction score*, defined as the degree to which learned representations in the trained model capture the full similarity structure of the environment abstracted across items, learning episodes, contexts and modalities (Jackson et al., 2021; McRae, Cree, Seidenberg, & McNorgan, 2005; Rogers & McClelland, 2004). Good conceptual representations should express similarity amongst items in the environment based on their full complement of properties, regardless of which properties are observed or inferred in a given situation. Whilst behaviour must be context-sensitive, such context-invariant, multimodal conceptual representations are critical to allow past events to inform semantic behaviour through generalisation across tasks, events and sensory modalities (Jackson et al., 2021; Lambon Ralph et al., 2017; Lambon Ralph & Patterson, 2008).

## Results

### Simulation 1: Effects of delayed semantic control with full connectivity from control

The first simulations used an architecture with full connectivity from the control units throughout the network (all three connections shown in Figure 1). This connectivity pattern was the starting point in the previous explorations, and while the presence of multiple long-range connections may not be biologically plausible, it obviates the need to rely on any assumptions about the precise connectivity pattern. These trainable weights allow the Control Layer to influence the flow of activation through the model at all layers. Precisely which units were influenced by control, and in what manner, was shaped solely by learning. The ANOVA showed that, in this model, the length of the maturational delay reliably influenced both conceptual abstraction (F(5, 395)=81.031, p<.001, η²=.788) and training time (F(3.468, 83.228)= 177.248, p<.001, η²=.881).

Neither the transition function (conceptual abstraction score; F(1, 79)=0.104, p=.748, η²=.001; training time; F(1, 24)=3.668, p=.067, η²=.133), the transition window (conceptual abstraction score; F(2, 158)=1.609, p=.203, η²=.040; training time; F(2, 48)=1.195, p=.312, η²=.047), nor their interactions explained significant additional variance in either metric, so the data were collapsed across these factors to further analyse the effects of maturational delay.

Figure 2.A. shows the distribution of conceptual abstraction scores across model runs at each level of maturational delay. Compared to the baseline condition (zero delay), a short maturational delay improved conceptual abstraction (at 1000 epochs; t(958)=2.758, p=.030, CI=.015, .003; at 2000 epochs; t(958)=4.792, p<.001, CI=.008, .020); an intermediate delay showed no difference (3000 epochs; t(958)=.867, p=1, CI=-.003, .009); and longer delays showed significantly worse scores (4000 epochs, t(958)=-5.430, p<.001, CI=-.023, -.011; 5000 epochs, t(958)=-13.015, p<.001, CI=-.048, -.036). These effect sizes were, however, relatively small except for the most extreme delay (for improvements at 1000 and 2000, d = 0.178 and 0.309 respectively; for decrements at 4000 and 5000, d = 0.351 and 0.840 respectively). Thus, increasing the maturational delay produced an inverted U-shaped effect on the quality of the model’s conceptual representations with modest initial improvement followed by modest reductions and significant degradation only at the longest lag.

**Figure 2.**
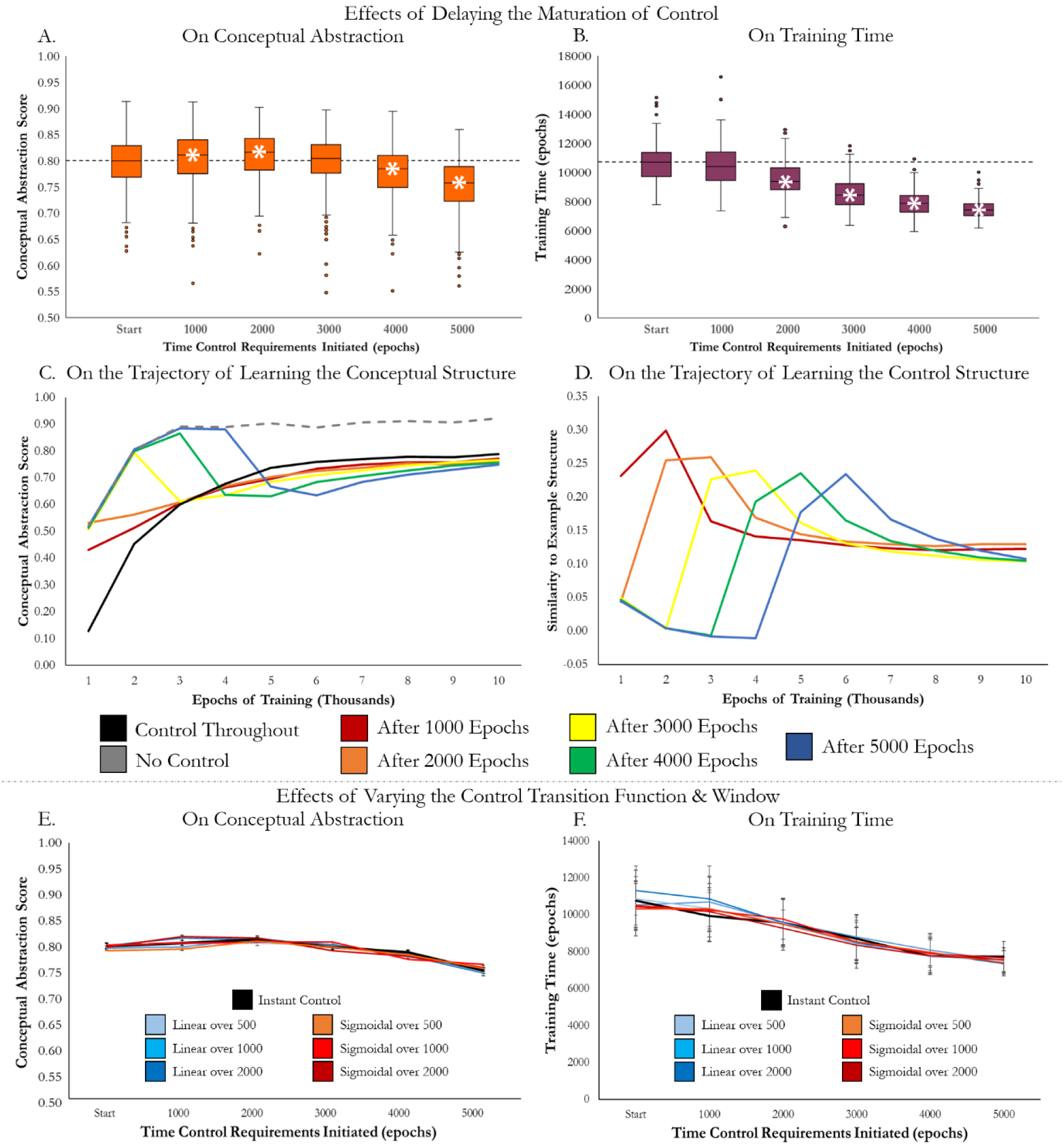
The effects of a maturational delay in learning control on conceptual abstraction, learning trajectory and training time. Control is added in full from the start, or after 1000, 2000, 3000, 4000 or 5000 epochs and the Control Layer has full connectivity to the rest of the model. Results are collapsed across the various protocols for gradually adding control. A. The effects of a maturational delay in control on the conceptual abstraction score. B. The effect of a maturational delay in control on the time taken to train the model. A&B. Bars signify the median value and the 25^th^ and 75^th^ percentile values. White asterisks signify a significant effect of the maturational delay in adding control in contrast to including control requirements from the start of training. Dashed lines allow easy comparison of each delay period to this baseline. C. The effects of a maturational delay in control on the evolution of the mean conceptual abstraction score across training. D. The effects of a maturational delay in control on the trajectory of the representation of the context signal, or control structure in the Hub Layer. C&D. The mean similarity of the representations in the Hub Layer to each structure is displayed after every 1000 epochs of training. Results for a model where control is never added is shown with a dashed line for comparison only as this model would never meet all core requirements of the human semantic system. A maturational delay results in greater representation of the conceptual than the contextual structure in the deep hub throughout training. In contrast, including control requirements throughout results in first learning the control structure in the deep hub before replacing it with the contextual structure. E&F. The effect of the developmental period without control on conceptual abstraction and training time are displayed for each of the different protocols and lengths for adding control (red = sigmoidal, blue = linear, lighter = control added over fewer epochs, darker = control added over more epochs). An additional instant protocol is shown for comparison (black). Error bars represent the standard error of the mean.

Figure 2.B. shows the distribution of training time across model runs at each maturational delay level. Here increasing maturational delay was unambiguously and monotonically associated with shorter training times relative to baseline (1000 epochs, t(298)=-1.349, p=.891, CI=-541.81, 101.06, d=.156; 2000 epochs, t(298)=-7.421, p<.001, CI=-1415.36, -822.02, d=.857; 3000 epochs; t(285.992)=-14.576, p<.001, CI=-2426.90, -1849.43, d=1.683; 4000 epochs; t(256.616)=-20.307, p<.001, CI=-3029.77, -2494.09, d=2.345; 5000 epochs; t(220.638)=-24.701, p<.001, CI=-3403.65, -2900.66, d=2.852). Based on these Cohen’s d values, a maturational delay of 2000 epochs or more produces a large or very large reduction of training time; indeed, a sufficiently long developmental period without control can reduce the training time by over a quarter. If the longest delay is avoided, a large reduction in training time may be gained whilst preserving the quality of the representations.

#### Interim Discussion

Both patterns are counterintuitive. Recall that conceptual abstraction measures how well the model’s learned representations capture a target conceptual structure, computed by considering all of an item’s properties tabulated across all modalities, regardless of which modalities are relevant for the current task context. Prior to the onset of semantic control, the model is trained to directly activate all such properties—thus one might expect that a longer maturational delay should produce monotonically *better* conceptual abstraction. Whilst small improvements in conceptual abstraction are possible with a short delay, increasing the delay further resulted in a *decrement* to conceptual abstraction. Overall, delaying semantic control only induced small changes in conceptual abstraction, except where very long delays resulted in a large decrement. With regard to training time, recall that the criterion for mature performance is based solely on the model’s ability to generate the correct output for the controlled, task-specific patterns. Thus, one might expect the fastest path to this objective would be to train the model with such patterns from the outset. Instead, this condition produced the *longest* training time. The more we delayed the introduction of controlled training patterns, the shorter the overall training time to master these patterns. The addition of a moderate delay (between one third and one half of the subsequent learning time) provides a substantial reduction in training time whilst maintaining the same or a slightly reduced level of conceptual abstraction. Thus, we find evidence for positive effects of a delay, particularly on the speed of learning.

To better understand these effects, we undertook some additional simulations and analyses. We first considered how the similarity structure of the hub representations evolved over time at each maturational lag. In each delay condition, we trained 80 independent models for 10,000 epochs (regardless of the performance criterion), stopping the training every 1,000 epochs and recording the patterns of activation evoked over the hub units for each item in every task. We then considered how the similarities amongst these learned representations correlated with two different target matrices: (a) the *conceptual similarity matrix* which includes the relationship between all features in each concept and is used to generate the conceptual abstraction score, and (b) a *contextual similarity matrix* that indicates the degree to which different tasks activate overlapping units in the Control Layer irrespective of the item being processed. Thus, the contextual similarity matrix expresses the relevance of the various input and output modalities across trials. The conceptual and contextual similarity matrices are uncorrelated with one another, so their individual correlations with the similarities expressed in model representations indicate how much model representations are influenced by conceptual versus task structure.

Figure 2.C. shows evolving correlations with the target conceptual similarity matrix as the model learns, for each delay condition and as a comparison, for the model trained without semantic control requirements (grey line). When control is present from the outset, correlations with conceptual structure increase gradually and monotonically. With delayed onset of semantic control, the model learns conceptual structure much more rapidly in initial epochs, but the introduction of controlled learning patterns significantly disrupts these trajectories, leading model representations to ultimately capture somewhat *less* conceptual structure with long delays than had control been present from the onset. This disruption is larger for longer maturational delays—the more the control-free model has mastered the conceptual structure, the greater the disruption to representations when control is introduced. All model representations recover somewhat from this disruption, with moderate lags achieving similar levels of conceptual abstraction to those with control throughout. However, those arising under very late-developing semantic control never “catch up” to acquire conceptual representations of comparable quality to the other models.

To understand why this disruption occurs, consider the most extreme delay, in which the model has learned good conceptual representations and is well on the way to producing the correct pattern of activation for each item across all three modalities—so given the input for *dog* in M1, true properties of *dog* in all modalities are activated and properties not true of *dog* are inactive. When a controlled pattern is introduced—say the model views the dog in M1 and must activate its properties in M2 but not M3—the units in task-relevant modalities (M1 and M2) are already near the correct state and will generate little error. Likewise, units in the irrelevant modality (M3) that are not true of dogs are near the correct state and generate little error. Thus, the error will mainly be driven by activation of units true of the dog but irrelevant to the task, and weight changes throughout the system will be driven to reduce the activation of these units.

Such changes could occur within the semantic representation network, or within the weights projecting from the active semantic control units. If the disruption of conceptual structure arises via input from control, then different instances of the same task (mapping from M1 to M2) should disrupt conceptual representations in a similar way, since all such instances are represented in the same way in the Control Layer. Conversely, instances of different tasks should disrupt the representations in different ways. Put differently, if the disruption to learned conceptual structure arises via inputs from control, the disrupted hub representations should begin to partially reflect the similarity structure of the task representations. Figure 2.D. shows just this pattern: as semantic control is introduced, representational structure in the hub increases its correlations with task structure, just as correlations with conceptual structure decline.

Correlations with task structure peak when the hub representations show the least amount of conceptual structure, and as conceptual structure recovers, the impact of task structure diminishes. This pattern suggests that some disruption of conceptual structure in the hub arises because control units directly impact the patterns of activation in the network, causing the network to represent a blend of conceptual and contextual structure, with continued learning promoting the conceptual structure.

Figure 3 shows the representational similarity structure in the hub at each time point across different lags. For reference, the bottom panels show the conceptual similarity (left), the contextual similarity (middle), and the similarities amongst output activations for all controlled training patterns (right). Where semantic control is present throughout training, the hub initially learns a roughly equal blend of the conceptual and contextual similarities, and over time, the contextual similarities gradually fade as the conceptual structure comes to dominate. Achieving a high level of conceptual abstraction relies on the functional specialisation of the hub for conceptual similarity, as distilling this crossmodal, cross-context structure is only possible in an area with connections to all sensory modalities. When control is delayed, the hub first encodes conceptual structure, and the longer the delay, the more structure it captures (i.e., similarity matrices increasingly resemble the context-independent multimodal conceptual structure at bottom left). The introduction of semantic control always produces a large initial effect, with task similarities very clearly apparent, changing the nature of the hub representations to reflect contextual similarity. Conceptual structure gradually re-emerges, yet changes later in learning are slow due to reductions in the effective learning rate with smaller error values. For shorter delays the conceptual structure in the hub is recovered to an equivalent level when the task criterion is reached, yet with longer delays conceptual abstraction never reaches the level observed when control is present from the start. Thus, control may be introduced after a delay to speed learning but to avoid detrimental effects on the representations learnt, control must be introduced before a critical point of learning.

**Figure 3.**
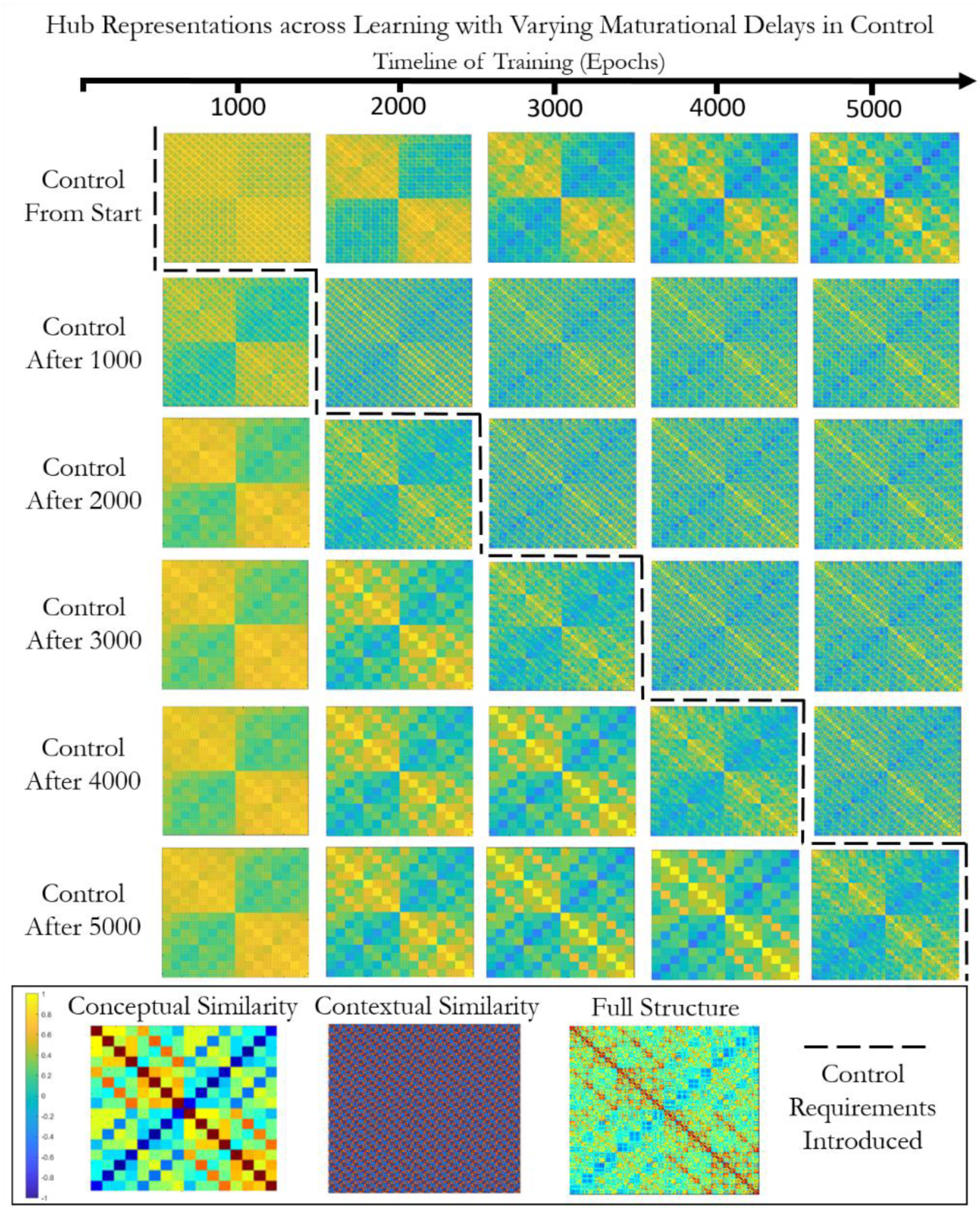
Visualising the model representations during learning across varying maturational delays in control. Control was present from the start of training or added around 1000, 2000, 3000, 4000 or 5000 epochs (dashed line). Control was added gradually using the sigmoidal protocol over 1000 epochs in the model with connectivity between the Control Layer and every other layer. Similarity matrices were constructed comparing activation in the Hub Layer across all the examples in each context. For comparison the target conceptual similarity matrix used to determine the conceptual abstraction score (i.e., the context-independent representation structure) is displayed, alongside the contextual similarity structure (reflecting the context signal only, which denotes the input and output modality of each trial) and the full structure (the combination of these contextual and conceptual structures). The Hub Layer specialises in the abstracted conceptual representation structure. Without control this functional specialisation is present in the hub from the start of training, whereas with control the context signal has a large initial effect on these deep representations.

These observations explain why long-delayed semantic control degrades conceptual abstraction in the hub. Initially one might expect degraded abstraction to produce longer learning times as well: if the model does a poorer job of representing cross-item, cross-modality structure, then learning should generalise less well across related times, leading to slower overall learning.

Yet we observed *faster* learning the longer control was delayed. To understand why such speeding may occur, consider again what happens in the longest delay condition: the model comes close to mastering all properties for all items across all modalities prior to the onset of semantic control. This in turn means that, for any given controlled pattern, the majority of output units will already be in the correct states: given a task where the network views *dog* in M1 and must generate output in M2 but not M3, all M1 and M2 units will be near the correct active or inactive state, but so will many units in M3—namely, all those that are not properties of dogs. If each item activates half the properties in each domain, then learning prior to the onset of control will place 5/6 of the output units in the correct state for any task that maps from one domain to another. Even for tasks involving only a single modality (such as retaining a visual stimulus in mind when it has vanished), all units in the relevant modality and half the units in the other modalities—or 2/3 of output units in total—will be in the correct state. The sparser an item’s properties are in each modality, the more units in irrelevant modalities will already be in the correct state as control trials are introduced. In other words, extensive experience with full patterns prior to control benefits performance for the majority of units in each controlled pattern. Note also that in the case of delayed control only, the initial learning in the hub reflects its ultimate role in representing conceptual similarity. With control present from the start, the model first learns to represent the contextual structure in the hub which then must be replaced by the conceptual structure. While some changes must occur when semantic control is introduced, they may be faster than this functional reorganisation across the representation network.

### Simulation 2: Effect of control connectivity

Simulation 1 showed that, when control connects to all layers of the semantic network, a moderate maturational delay to semantic control onset can greatly speed training time, without large differences in conceptual abstraction. Alternatively, a short delay could modestly improve conceptual abstraction. However, full connectivity from control to each layer of the representation network is neither biologically plausible (due to metabolic and packing constraints on long-range connections in the brain) or beneficial for performance. Previous analyses comparing the effect of the locus of the connection from control in this model, have demonstrated the importance of avoiding direct connections between control and the hub for successful conceptual abstraction (while connections to either more shallow layer resulted in equivalent performance) (Jackson et al., 2021). This could explain the apparent lack of structural connections between the PFC and ventral ATL in tractography assessments (Binney, Parker, & Lambon Ralph, 2012; Jung, Cloutman, Binney, & Lambon Ralph, 2016). However, these findings may differ if semantic control is introduced later.

Therefore, in Simulation 2 we considered whether the benefits of delayed semantic control depend upon the locus of the connection from control to answer three critical questions i) would connections restricted to shallower layers show the same pattern of changes with a maturational delay, ii), would the delay obviate the need to avoid a direct connection from control to the hub and iii) does the maturational delay otherwise change the most advantageous connectivity pattern? Simulation 2 replicated simulation 1 but restricted connections from the Control Layer so that they projected only to (1) the spokes, (2) the intermediate hidden layer, or (3) the hub.

Additionally, since the transition function and width of the transition window had no effect in simulation 1, we employed only a single transition protocol involving a sigmoidal function extended over 1000 epochs. 80 models were trained for each architecture at each maturational delay. As before we measured the influence of these factors on the model’s conceptual abstraction score at the end of learning, and on the total training time.

Results are shown in Figure 4. Conceptual abstraction scores were influenced by the location of the control connection (F(2, 158)=3414.917, p<.001, η²=.977), the maturational delay (F(5, 395)=14.449, p<.001, η²=.155) and their interaction (F(10, 790)=9.806, p<.001, η²=.110). Conceptual abstraction was always significantly worse when control connected to the Hub versus the Intermediate (t(822.103)=-56.164, p<.001, CI=-.228, -.213, d=3.989) or Spokes Layers (t(746.774)=-61.795, p<.001, CI=-.240, -.225, d=3.99), replicating Jackson et al., (2021). When control connected to the hub, maturational delay improved conceptual abstraction (at 1000 epochs; t(158)=2.989, p=.016, CI=.012, .057, d=.473; 2000 epochs; t(158)=5.187, p<.001, CI=.037, .083, d=.820; 3000 epochs; t(158)=5.761, p<.001, CI=.042, .086, d=.911; 4000 epochs; t(158)=2.895, p=.022, CI=.011, .057, d=.458; 5000 epochs; t(158)=3.381, p=.005, CI=.016, .059, d=.535), but never produced scores comparable to other connection conditions due to the large negative effects of the connection locus. Thus, even with maturational delay, conceptual abstraction is best promoted by architectures that prevent the direct impact of control on the hub. This may explain the lack of structural connections identified between the PFC control site and the multimodal vATL hub in the cortical semantic system (Binney et al., 2012; Jung et al., 2016).

**Figure 4.**
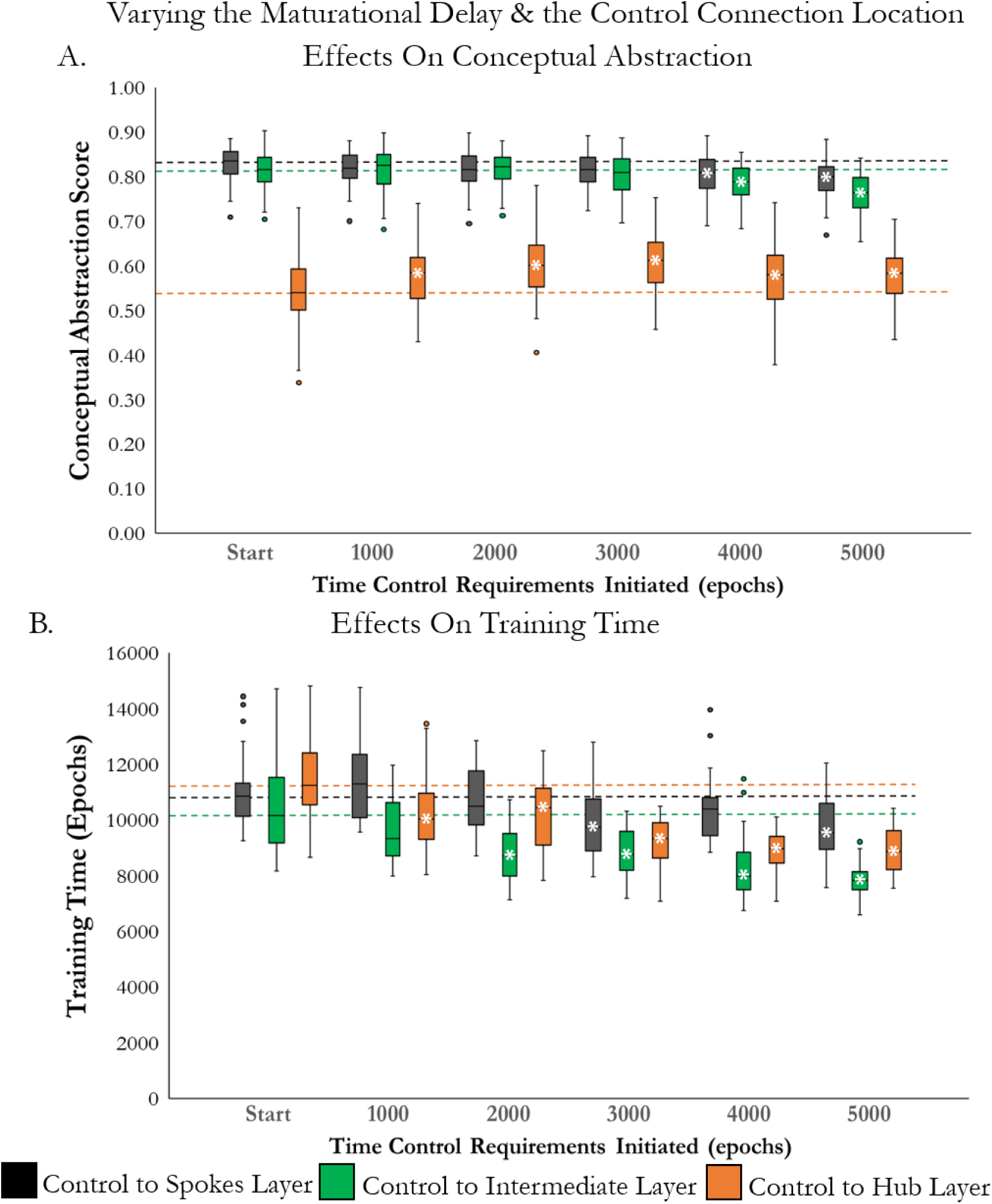
Assessing the effects of a maturational delay in learning control on conceptual abstraction score and training time with control connections to the different layers. The Control Layer is connected to the Spokes Layer (black), Intermediate Layer (green) or Hub Layer (orange). Control is present from the start, or added gradually around 1000, 2000, 3000, 4000 or 5000 epochs (with a sigmoidal protocol over 1000 epochs). A. The effect on conceptual abstraction. B. The effect on training time. Bars signify the median value and the 25^th^ and 75^th^ percentile values. White asterisks signify a significant effect of the delay in adding control in contrast to the inclusion of control requirements from the start of training. Dashed lines allow easy comparison of each delay period to the baseline level where control is present from the start.

Control connections to the Spoke versus Intermediate Layer produced reliably better conceptual abstraction (t(934.139)=4.265, p<.001, CI=.006, .018) but the effect size was small (d=.275). Models with either connection locus showed equally good conceptual abstraction for delays up to 3000 epochs (Spokes Layer connection; at 1000 epochs; t(158)=-1.868, p=.318, CI=-.022, .001; at 2000 epochs; t(158)=-1.993, p=.240, CI=-.024, .000; at 3000 epochs; t(158)=-2.488, p=.069, CI=-.026, -.003; Intermediate Layer connection; at 1000 epochs; t(158)=-0.134, p=1, CI=-.014, .013; at 2000 epochs; t(158)=0.163, p=1, CI=.011, .013; at 3000 epochs; t(158)=-1.460, p=.731, CI=-.022, -.003). As with simulation 1, longer lags produced reliable decrements in both cases, with only the longest lag producing a large effect size (contrast to baseline for connections to the Intermediate Layer: at 4000 epochs; t(158)=-4.886, p<.001, CI=-.043, -.018, d=.773; at 5000 epochs; t(158)=-8.021, p<.001, CI=-.068, -.041, d=1.268; for connections to the Spokes Layer: at 4000 epochs; t(158)=-3.616, p=.002, CI=-.035, -.010, d=.572; at 5000 epochs; t(158)=-5.369, p<.001, CI=-.045, -.021, d=.849). Thus, in both cases the effects were similar to those observed in simulation 1 with large changes in conceptual abstraction resulting from the longest delays only. However, in neither case did maturational delay produce the modest positive effects and resulting inverted-U shape observed in simulation 1.

Training time also varied with control connection (F(2, 48)=40.169, p<.001, η²=.626), maturational delay (F(5, 120)=32.659, p<.001, η²=.576), and their interaction (F(6.153, 147.668)=3.612, p=.002, η²=.131; see Figure 4.B.). Across all connection loci later maturation of control sped learning (for each contrast to baseline t(298)>4.2, p<0.001). However, the extent of the speeding was greater for the deeper connections (Hub Layer maximum d=2.221; Intermediate Layer maximum d=2.139) than the Spokes Layer (maximum d=1.106). As the connection to the Intermediate Layer also started with lower training times than the control projection to the hub, the fastest training occurred when control projected to the Intermediate Layer (contrast to Spokes Layer connection t(298)=-9.958, p<.001, CI=-1882.411, -1261.134, d=1.150; contrast to Hub Layer connection t(298)=-5.466, p<.001, CI=-1210.621, -569.632, d=.631). At all lags, a control connection to the Intermediate Layer resulted in the fastest training. With this connection locus, as in Simulation 1, moderate and long maturational delays decreased training times substantially (at 1000 epochs; t(48)=-1.901, p=.316, CI=-1494.258, 41.778, d=.316; at 2000 epochs; t(48)=-4.268, p<.001, CI=-2313.239, -831.721, d=1.207; at 3000 epochs; t(38.141)=-4.369, p<.001, CI=-2267.188, -831.532, d=1.236; at 4000 epochs; t(48)=-5.252, p<.001, CI=-2821.033, -1258.97, d=1.485; at 5000 epochs; t(31.551)=-7.563, p<.001, CI=-3177.669, -1842.971, d=2.139). Thus, with an intermediate control connection locus, a moderate maturational delay (3000-4000 epochs or around half the subsequent training time) results in greatly speeded training in the context of relatively preserved conceptual abstraction.

#### Interim Discussion

Simulation 2 provides the answers to our three key questions. First, it is clear that a maturational delay in the introduction of semantic control cannot obviate the need to avoid direct connections from control to the multimodal hub. As in prior work, direct connections from control to the hub drastically impair its ability to uncover the latent conceptual structure across contexts and modalities even with the positive effects of a maturational delay. While the delay does improve conceptual abstraction, the improvements provided by even the longest delays are much smaller than the negative effects of this direct connection. Thus, control regions should not directly connect into the multimodal hub. As noted in Jackson et al., (2021), this conclusion from simulation data accords well with the anatomical structure of the cortical semantic network, as there is no evidence for direct structural connections between PFC regions critical for semantic control and the multimodal vATL hub (Binney et al., 2012; Jung et al., 2016).

Second, models that respect this architectural constraint replicate the critical findings from the fully connected model: a maturational delay in the development of semantic control produces greatly speeded learning whilst maintaining conceptual abstraction ability. Thus, the late onset of semantic control has positive effects even with a more plausible restricted connectivity pattern that avoids the detrimental connection from control to the hub. These models do not show improvements in conceptual abstraction ability with a delay unlike Simulation 1. When the absence of these direct connections allows the semantic hub to remain insulated from control inputs, maturational delay cannot improve conceptual abstraction further by protecting the hub from control signals early in learning. Thus, it would be advantageous to have both a connection from control to semantic representation regions closer to sensory input (to preserve conceptual abstraction) and a moderate maturational delay in semantic control (to speed learning). Maturational delay and system connectivity can conspire to promote faster overall learning with high levels of conceptual abstraction.

Third, by considering the effect of a developmental delay we are able to separate the consequences of a connection to the two shallow layers. Whilst the effects of connecting to Intermediate versus Spokes layers appear similar when control is presented throughout training, the large improvements in speed were greater when control connected to the Intermediate Layer. Thus, the present results highlight the benefit of both a maturational delay in semantic control and a connection from control to regions located intermediately between sensory input and multimodal hub regions. Whilst long delays result in compromised conceptual abstraction ability, moderate delays can have very large speeding effects with either fully or relatively preserved conceptual abstraction. The optimal length of the delay is around one third to one half of the subsequent learning time, depending whether some trade-off with conceptual abstraction is considered advantageous.

#### Part 2: Meta-analysis of Developmental Data

The simulations suggest that a delay in the maturation of the semantic control system can lead to overall faster mastery of controlled semantic behaviours and to better structured semantic representations. Does the literature contain evidence of such a delay? As noted in the Introduction, research on the development of semantic control is lacking, with much of the prior control research tapping into distinct domain-general executive control processes and/or utilising tasks lacking the retrieval of any semantic knowledge. Thus, it is unknown whether semantic control develops later than semantic representation. Next, we assess existing research at the border of executive control and semantic cognition to determine whether it can provide any clues as to the development of semantic control.

Some traditional executive tasks do require access to semantic knowledge. However, canonical measures typically involve asking the child to generate behaviours in direct opposition to the common meaning of the stimulus. For instance, the child might be instructed to say “night” to a picture of a sun and “day” to a picture of the moon (see also Grass/Snow, Less is More, the Hand game, Reverse Categorisation) (Gerstadt C.L., Hong Y.J., & A., 1994). Rather than having the child generate a correct, context-appropriate and ecologically realistic response to the stimulus, these tasks instead require complete reversal of the correct inference to behave in accordance with an arbitrary rule that has little connection to how knowledge is used in daily life. Indeed, of 24 common executive tasks surveyed by Carlson et al., (2005), none have the properties that characterise semantic control: retrieval of true, non-salient but context-appropriate semantic information about a perceived stimulus.

A potential exception arises in the first (and often unanalysed) block of the gold-standard test of executive development, the dimensional-change card-sorting (DCCS) task, a child-appropriate variant of the Wisconsin card sort task (Zelazo, 2006). Children are instructed to match cards depicting animals and objects to a standard based either on colour or “shape,” that is, the category to which the depicted item belongs (cow, boat, car, etc.). After a series of trials with one instruction (e.g., “Let’s play the colour game. Which one is the same colour?”), the instructor switches the matching criterion (e.g., “Now let’s play the shape game. Which one has the same shape?”). Executive control is measured as the ability to quickly shift behaviour to accord with the new instruction. Children typically perseverate on whichever rule was first used until about age 4 years, consistent with the developmental timeline for the onset of inhibitory executive control across domains. However, it is not the task-switching that is central to semantic control, but the ability to perform this task at all. It is the behaviour in the first, *pre-switch* block of the DCCS that is of greater interest for understanding development of semantic control. The “shape” condition of the task clearly requires semantic control: given the same stimulus, the child must make different matching decisions depending upon the instruction context, and matching by “kind of thing” arguably requires semantic categorisation of the item depicted on the card and on the standard. Early work with this task suggested that children younger than 3 years were unable to follow the matching rule despite knowing the correct answer (Zelazo & Reznick, 1991) but more recent work employing simpler touchscreen methods have found that children as young as 2 years (the youngest age assessed) reliably choose the match that accords with the instruction in the first test block, and indeed succeed at rates comparable to children at 3-3.5 years of age (Blakey, Visser, & Carroll, 2016). Such results suggest that children can successfully shape their behaviour in a context-appropriate manner as young as 2 years, much earlier than the canonical onset of full inhibitory executive control.

Similarly, studies that have focused directly on semantic control in language comprehension report successful performance at the youngest ages tested. For instance, Deak (2000) taught children aged 3-6 years an unfamiliar word (e.g., “rebek”), varying the predicate used to introduce the item. All children were shown the same novel reference object, but some were told it “looks like a rebek”, some that is was “made of rebek”, and some that it “has a rebek.” They then asked children which of several options also looked like, was made of, or had the property. Even 3-year-old children shaped their generalisation behaviours differently depending on the predicate context—that is, they showed clear evidence of semantic control, exhibiting different choice behaviours based on the predicate context. Such results align well with classic findings of inductive projection over development showing different generalisation profiles for different kinds of newly-learned information in children as young as 3 years (e.g., Carey, 1987), again suggesting that such semantic control may develop earlier than executive functioning writ large.

However, critical questions about when a given cognitive faculty develops are difficult to answer broadly because successful performance on any given metric can depend on other demands of the particular task, such as the complexity of the required motor behaviours, or the nature / familiarity of the stimulus materials. For instance, 3-year-olds who fail to place cards correctly on switch blocks of the DCCS can often succeed in pointing to the correct location; children who succeed in conforming to the new response rule in DCCS at age 4 years can nevertheless fail to do so in parallel tasks involving emotional inference in speech, etc. (e.g., Morton & Munakata, 2002). These observations suggest that the developmental timeline for semantic control remains unestablished. Critically, the core prediction of the simulations—that children should pass through a phase during which they possess relevant semantic knowledge for a given task but cannot control retrieval to generate context-appropriate responses—remains untested. Furthermore, any such test requires a task that can be conducted with children as young as 2 years and that can assess both whether the child possesses the target knowledge and whether they can control retrieval of the knowledge in a context-appropriate manner.

In this section, we consider research arising from an early controversy in conceptual development that makes extensive use of such a task. The controversy arose from Inhelder and Piaget’s (1958) early view that children’s conceptual representations first capture thematic associations that eventually give way to taxonomic relationships, and the subsequent competing view that early concepts primarily reflect taxonomic structure at either basic (Rosch et al., 1976) or even more general or abstract (Mandler & McDonough, 1993) levels. To adjudicate these possibilities, many studies have employed a task now known as *triadic match-to-sample*: the participant must decide which of two items best matches a third sample item. For instance, a participant might judge which “goes better with” a picture of a horse: a saddle or a deer? Choice of the saddle indicates a thematic match, since horses and saddles occur together in many scenarios but are distinct kinds of things. Choice of the deer indicates a taxonomic match, since horses and deer don’t often occur together in familiar scenarios but are similar kinds of things (i.e., four-legged land mammals).

Throughout the 1970’s and 80’s many such studies were conducted, with some favouring the view that taxonomic knowledge trumped thematic knowledge and others suggesting the reverse (e.g., Bauer & Mandler, 1989; N. W. Denney, 1974; Greenfield & Scott, 1986; Smiley & Brown, 1979). Eventually it became clear that the results depend critically upon the wording of the instructions (Estes et al., 2011; Lin & Murphy, 2001; Waxman & Namy, 1997)—that is, upon the task context. If asked to choose which item has the same name, children most often selected a taxonomic match. If asked which item “goes better with” the target, they more often selected the thematic match. In 2001, Lin and Murphy conducted a large triadic matching study with adult participants, assessing the effects of many different instructional contexts. The results revealed a gradation of choice preferences, with some contexts strongly eliciting taxonomic choices (e.g., “called by the same name”), others reliably eliciting thematic choices (“goes with”), and still others yielding somewhat ambiguous results.

These findings indicate how the triadic matching task indexes semantic control when the choice options include both a taxonomic and thematic match. In such cases the semantic representation system yields information about precisely how each option matches the target, and the participant must use some representation of the current goal to select which kind of information is most relevant to the task. The instructional cue is a form of context, providing this goal representation. Successful semantic control is signalled by the ability to reliably choose the taxonomic option under one instructional context and the thematic option under another.

Suppose, however, that a participant fails this task by choosing unsystematically. Such a failure might arise (a) because the participant is unable to use the control system to choose the correct kind of semantic information, or (b) because the participant does not possess the semantic knowledge needed to choose correctly. These possibilities can be adjudicated by assessing whether the participant correctly matches a given item for triads where the second option is *unrelated* to the sample. If the participant is able to choose the correct match in the context of unrelated distractors, this indicates that they possess sufficient knowledge of the relevant semantic information to find the target under a given set of instructions. That is, selecting the match when the target appears with an unrelated item does not strongly tax semantic control, but does assess knowledge of the relevant matching property. Thus, a signature of good semantic knowledge representations with underdeveloped semantic control would arise when a participant succeeds at a given kind of matching when the second option is unrelated to the target, but fails when the second option is related to the target in a manner irrelevant to the task context.

With these ideas in mind, we undertook a meta-analysis of the developmental literature on triadic matching for taxonomic and thematic knowledge, aiming to assess whether knowledge of taxonomic and thematic relations emerges prior to, or in parallel with, semantic control. We identified studies which utilised a match-to-sample task with either thematic or superordinate taxonomic matching or both and reported the frequency of choosing the taxonomic or the thematic match.

Studies were designated as either low or high control. In l*ow control studies* the target matched the sample either taxonomically or thematically, and remaining options were semantically unrelated to the sample. These trials thus indicated whether the participant possessed taxonomic or thematic knowledge sufficient to choose the target without having to select what kind of semantic information was relevant for the task. In *high control studies* all trials have both a thematic and taxonomic match, but the instructional context indicates which kind of information should drive the selection. These trials thus require both the relevant semantic knowledge and the ability to select the correct kind of information for the task.

Note that taxonomic and thematic relationships are encoded by different varieties of environmental structure—for instance, dogs, wolves and foxes are similar by virtue of shared parts, behaviours, and appearances, but dogs, bones, and leads are similar by virtue of common co-occurrences with one another and with other contextual cues (e.g., Binder et al., 2016).

Consequently, selecting the taxonomic versus thematic relationships relies upon selecting some set of relevant features and inhibiting a different set of irrelevant features, whether these are represented in the same or different brain areas. This is the same operation performed by the computational models in Part 1.

For the low control studies, we did not further sort based on instructions as any semantic relationship cue should lead participants to choose the correct (taxonomic or thematic) target over an unrelated concept. For the high control studies, we analysed the instructions provided to participants and selected only those previously identified as reliably promoting a thematic or taxonomic decision in adults (D. R. Denney & Moulton, 1976; Estes et al., 2011; Greenfield & Scott, 1986; Lin & Murphy, 2001; Markman & Hutchinson, 1984). Thematic contexts were those cued by ‘goes with’, ‘goes best with’ or ‘goes together’ instructions, which imply concepts are found together in time or space. Taxonomic decisions were cued by the provision of a novel label for the standard (e.g., ‘this is a dax’) and the request to find another item in this category (e.g., ‘find another dax’). The novel word acts as a basic noun which typically relates to the taxonomic status of the concept (referred to as the noun-category bias; Waxman, Senghas, & Benveniste, 1997).

## Results

Figure 5 shows the mean proportion of taxonomic (top end of y axis) or thematic (bottom end) choices at each age and for each experiment type. To determine whether each age group had evidence of a conceptual representation structure, the proportion of correct responses in the taxonomic-only and thematic-only conditions were compared to chance (0.5), using one-tailed one-sampled t-tests. Studies with children as young as one showed chance behaviour, but by age two children chose the correct taxonomic or thematic match reliably more often than chance, so long as other options were semantically unrelated to the sample (taxonomic; t(3)=5.207, p=.007; thematic; t(3)=2.98, p=.029). When choices contained both a taxonomic and thematic match, two-year-olds chose equivalently (and near chance) regardless of the instructional context (taxonomic vs. thematic cues; t(3)=2.97, p=.393). Thus, by age two children appear to possess knowledge of both taxonomic and thematic relations, but are unable to use instructional context to determine the correct basis for matching. For ages 3 and 4, there were significant differences in responses to the two cue types, demonstrating the presence of some context-sensitivity (taxonomic vs. thematic cues; age 3; t(13)=2.222, p=.045; age 4; t(17)=3.985, p<.001). It is not clear whether these children can utilise both the taxonomic and thematic cues, or whether the (stronger) taxonomic cue is driving this difference alone (whilst thematically cued responding remains unsystematic). However, it is clear that this context-sensitivity is not yet at adult levels (taxonomic vs. thematic cues; age 5; t(5)=3.38, p=.01; age 6; t(1)=23.286, p=.014; age 7; t(1)=26.353, p=.024; adult; t(10)=11.972, p<.001). Context-sensitivity increases in strength throughout development (age 3; g=1.082; age 4; g=1.748, age 5; g=2.171; adult; g=7.364) and from age 5 onward children also display a preference for the thematic response when presented with a thematic cue. Context-sensitivity shows a large increase and approaches adult levels around the age of 6. Thus, consistent with the model predictions, children acquire the relevant semantic representations and can deploy them to perform the matching task as young as 2, but cannot begin to make systematic use of contextual information to guide task-appropriate semantic choices until age 3, with large increases in semantic control around 6 years.

**Figure 5.**
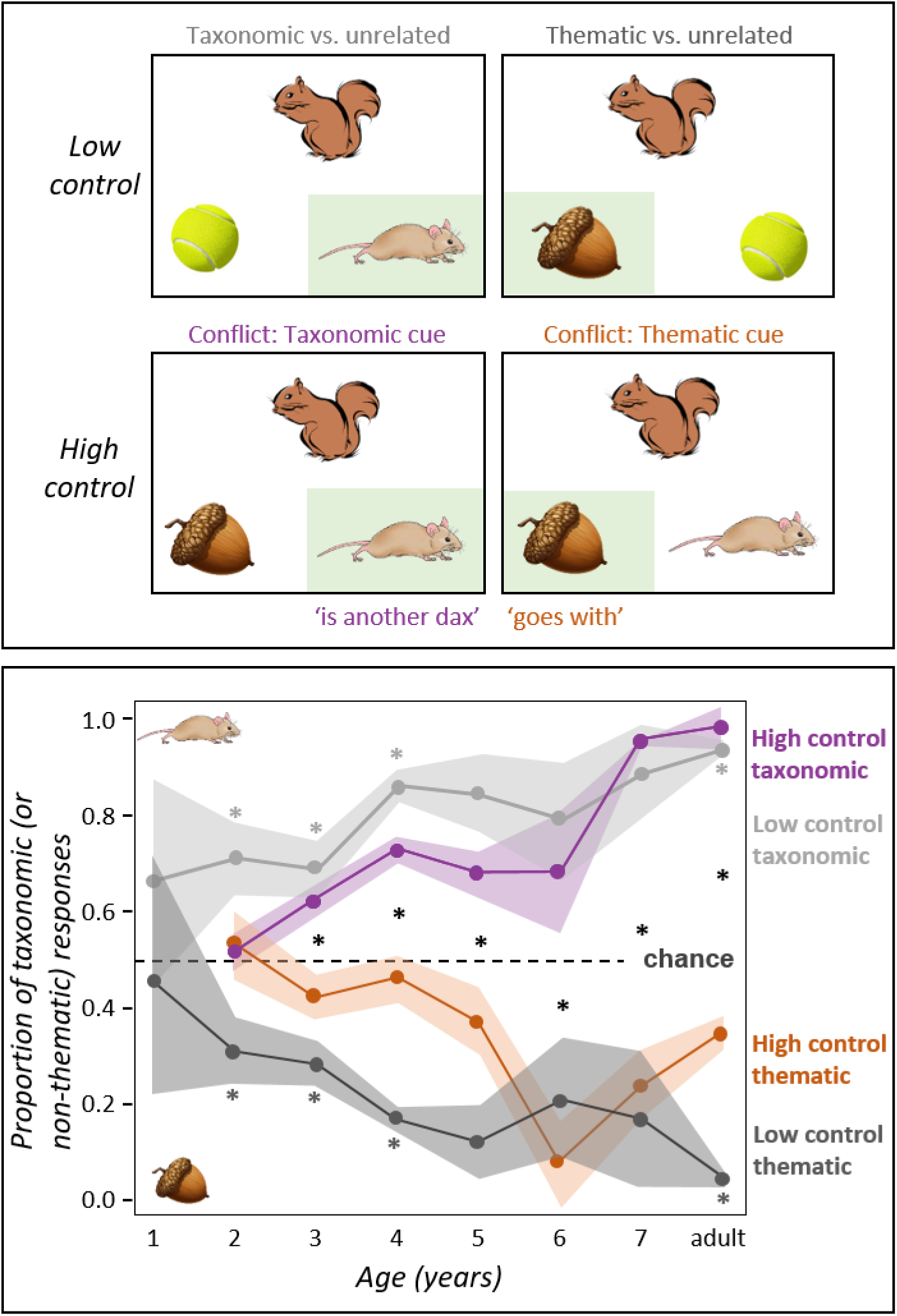
The emergence of semantic control after semantic representation in our meta-analysis of triadic comparison. Top panel: Examples of a match-to-sample task illustrating the two low control conditions (taxonomic vs. unrelated and thematic vs. unrelated concepts) and the two high control conditions (both including a taxonomic and a thematic match but differing in the instructional cue). The correct answers are highlighted in green. Bottom panel: Mean and bootstrapped confidence intervals indicating the proportion of times participants in different age groups chose the taxonomic (high proportion) or thematic (low proportion) target in each study type. Knowledge of taxonomic or thematic relationships is indicated by non-chance responding in the low control conditions (light grey and dark grey lines respectively, light or dark grey asterisks denote responding significantly better than chance, not all contrasts are possible). The ability to use the instructional cue to inform the choice of an appropriate match in high control conditions is indicated by the difference between the proportion of taxonomic versus thematic responding for the taxonomic (purple line) and thematic (orange line) cues. Significant context-sensitive effects indexing semantic control ability, are signified by the black asterisks.

## General Discussion

Building on prior suggestions (Chrysikou et al., 2011; Newport, 1990; Thompson-Schill et al., 2009), we considered whether the late maturation of semantic control might benefit development of controlled semantic cognition. Using a neural network architecture shown in prior work to promote both conceptual representation and semantic control, we assessed the effects of delayed semantic control in scenarios where (1) control regions connect to the entire semantic representation network, or (2) control connects only to specific levels of the network. In both cases, maturational delay of semantic control relative to semantic representation promoted conceptual development by reducing the time taken to learn the full controlled behaviour with little cost to representational structure except at the very longest delays. Modest improvements in conceptual abstraction ability were present after short delays with connections from control to the hub (including the biologically implausible full connectivity condition).

However, the positive effects of the delay were insufficient to negate the poor-quality representations acquired when control connected to the multimodal hub. Thus, anatomical connectivity that insulates the hub from the direct effects of control promotes good learning of conceptual structure even when semantic control is substantially delayed.

This finding accords well with studies of structural connectivity in the human semantic system. White matter tractography suggests *an absence* of direct structural connections between the PFC and the ventral ATL multimodal hub (Binney, Embleton, Jefferies, Parker, & Lambon Ralph, 2010; Chang, Halai, & Lambon Ralph, 2023; Jung et al., 2016). While false negatives are quite possible in tractography, this contrasts with the multimethodological evidence documenting connectivity between the PFC and a more distal area, intermediate between the sensory regions and the multimodal hub. Specifically, the posterior temporal cortex is well connected to both the ventrolateral PFC region important for semantic control and to the vATL hub (Binney et al., 2010; Chang et al., 2023; Jung et al., 2016; Matsumoto et al., 2004; Souter et al., 2022). Damage to (or inhibitory stimulation of) this posterior temporal site, the PFC itself, or the connection between these regions, results in impaired semantic control, suggesting the impact of control on semantic representation areas may occur via the interaction of PFC with posterior temporal cortex (Jefferies, 2013; Jefferies & Lambon Ralph, 2006; Souter et al., 2022; Whitney, Kirk, O’Sullivan, Lambon Ralph, & Jefferies, 2011). This accords well with the result that the fastest training was observed with both a maturational delay and control connected to intermediate model layers. The simulations thus suggest that delayed onset of semantic control, combined with an architecture that avoids direct connectivity from control to the multimodal hub, leads to substantially faster mastery of controlled semantic cognition, and consequently that a slow maturation of semantic control combined with anatomical constraints may be adaptive in this regard.

Consistent with this view, a new meta-analysis of the classic triadic match-to-sample task showed that a child’s ability to use context to retrieve a particular kind of semantic information—that is, their ability to control semantic retrieval—arises somewhat later than does acquisition of the semantic information itself. While children as young as 2 years old possess taxonomic and thematic semantic information sufficient to choose a correct match from among unrelated distractors, they do not have the semantic control processes required to produce context-sensitive responses when taxonomic and thematic matches compete for selection. Subtle but reliable context-sensitivity arises around age 3 years, earlier than the domain-general inhibitory control measured by most studies of executive development, and increases with age, with reliable use of both instructional cues occurring around 6 years. These data thus suggest that maturation of semantic control lags behind acquisition of conceptual knowledge and representations, as predicted by the models.

### Why does maturational delay speed learning?

The simulations suggest two reasons why delayed onset of semantic control may ultimately speed mature semantic cognition. First, prior to the onset of semantic control, each learning experience can activate all known features of a concept, whereas with control, each experience activates only a subset of features. Training without semantic control thus effectively increases the amount of exposure to each feature, producing faster learning and good approximation of cross-modal representational structure. Second, in allowing the different features of a concept to co-activate regardless of the current task, a period with limited semantic control can help the system learn to use highly similar conceptual representations across tasks, rather than learning distinct mappings in different contexts. That is, delayed semantic control may avoid a key challenge for controlled semantic cognition, namely the need to extract context-invariant knowledge whilst simultaneously producing context-sensitive responding. Instead, context-invariant representations may be acquired before there is a strong requirement for context-sensitive behaviour and the presence of a substantial “base” conceptual representation may then allow the continued use of shared representations across different tasks later in development.

While it may seem tempting to consider what the simulations suggest about the optimal length of delayed semantic control in cognitive development, several factors make the extrapolation non-trivial. One important limiting factor is the nature of the model training environment, which sacrifices ecological validity in favour of the strict control that allows us to measure conceptual coherence and speed of learning. The results make it clear that later development of semantic control is beneficial overall, but connecting this observation to real developmental trajectories will require models with richer and more realistic training corpora such as those recently studied by Giallanza et al. (2025) in related work. It will also require better assessment tools for tracking the true developmental profile of semantic control from ages 2-10, and for differentiating this profile from executive function more broadly.

Importantly, these effects depend upon the anatomical structure of the overall network. The beneficial effect of delayed semantic control on learning speed is greatest when control representations connect to deep or intermediate network layers, whilst acquisition of conceptual structure depends upon insulating the deep hub from direct influence of control. These observed differences cannot be simply due to the amount of experience with each feature, since this is matched across different control connections. Thus, anatomy and maturational delay can conspire to promote fast learning with good conceptual abstraction, with the maturational delay allowing the full network to lay down base representations that approximate the cross-context structure of the environment before control begins to shape behaviour and control interfacing at an intermediate part of the semantic representation network leaving the multimodal hub relatively insulated from control. The current analyses demonstrate this in the context of a relatively simple model environment, allowing explicit assessment of representation quality. Again, future work should assess the scalability of such assessments to the complexity of a real-world environment and the impact of selecting different features within the same versus different modalities.

### Implications for the development of controlled semantic cognition

Much prior work modelling conceptual development has focused on the emergence of representational structure rather than semantic control. This research has been successful in explaining how sensitivity to multimodal environmental structure can explain a variety of developmental phenomena, such as the progressive differentiation of conceptual knowledge (Mandler & McDonough, 1993; Poulin-Dubois & Pauen, 2017; Rogers & McClelland, 2004; Saxe, McClelland, & Ganguli, 2019), the primacy of basic-level naming (Rogers & McClelland, 2004; Rogers & Patterson, 2007; Rosch et al., 1976), temporary overgeneralisation of frequent object names and properties (Mervis, 1987), and illusory correlations (Chapman, 1967; Rogers & McClelland, 2004). Indeed, the considerable success of this viewpoint may indicate that conceptual development proceeds without the disruptive influence of semantic control. That is, the importance of semantic control for conceptual development may have been neglected precisely because its onset is delayed until much of the core conceptual structure is learnt, as suggested by our meta-analysis of triadic match-to-sample data.

Our results also suggest a new perspective on the development of taxonomic and thematic concepts—specifically, that patterns of developmental change reflect growing sensitivity to contextual information maintained by the maturing semantic control system rather than an overall bias toward learning or deploying a particular kind of semantic information at a given age. Indeed, changes in the salience of or bias toward taxonomic versus thematic knowledge cannot explain the full pattern of results, since our meta-analysis showed that, across ages, children are equally able to exploit either kind of information when distractors are unrelated to the target (and thus there is little need for semantic control). Instead, what changes across development is children’s increasing sensitivity to task context. In this sense taxonomic and thematic relationships may be more similar than is often considered, with both being encoded within the same system of representation (Jackson, Hoffman, Pobric, & Lambon Ralph, 2015; Lambon Ralph et al., 2017) rather than in distinct systems as some have proposed (e.g., Mirman, 2017; Schwartz et al., 2011), and with semantic control governing which contributes to behaviour in a given task. Our proposal is also consistent with prior work showing the difficulty that both children (Blaye et al., 2006; Blaye & Jacques, 2009; Mirman, 2017) and healthy adults (Landrigan & Mirman, 2018; Sachs et al., 2008) have in switching between taxonomic and thematic judgments, a task that requires shifting the task representation and hence the semantic criterion of a decision in real time.

### Relation to other changes in development that may be attributed to semantic control

Semantic control may be differentiated from other forms of executive control by emphasising that it 1) requires the use of context to shape the production of true and situation-appropriate information, and 2) applies this control over the access and manipulation of meaningful semantic content. The triadic match-to-sample task is of interest because it assesses this form of control without also placing other executive demands, such as task switching, inhibition of highly salient or prior responses, or temporal sequencing. Through that lens, semantic control begins to emerge around age three and improves through age 6. This general timeline appears consistent with broader studies of developmental changes that may be underpinned by emerging semantic control processes, alongside other changes in domain-general executive control or knowledge acquisition (Davidson et al., 2006; A. Diamond & Doar, 1989).

Whilst children under 5 years old perseverate on subcategories in semantic fluency tasks, older children can switch to a new subcategory when explicitly cued and then learn to employ this as an endogenous strategy (Snyder & Munakata, 2010). Around 5 to 6 years of age, children become sensitive to the presentation of stimuli in a set (Snyder & Munakata, 2013), and learn to switch flexibly between multiple meanings of a word (Blaye et al., 2006; Blaye & Jacques, 2009). Other work documents a qualitative change in semantic cognition around 6 years in which children cease to simply name objects and start to describe them (Funnell, Hughes, & Woodcock, 2006), perhaps reflecting flexible, context-sensitive behaviour. Research in the development of creativity provides another source of evidence, as such work often uses similar tasks to those employed in studies of semantic control (e.g. generating atypical uses of an object) and engages similar brain areas (Chan, Lambon Ralph, & Robinson, 2025; Cogdell-Brooke, Sowden, Violante, & Thompson, 2020; Jackson, 2021; Krieger-Redwood et al., 2025; Noonan, Jefferies, Visser, & Lambon Ralph, 2013). Children under 6 years old lack ‘functional fixedness’, performing better than older children or adults on tasks which typically lead to fixation on a particular use of an object in adults. For instance, after a demonstration where a box is used as a container, adults and older children are less likely to solve a problem requiring its use as a step than are younger children (German & Defeyter, 2000). Similarly, 5-year-olds outperform 7-year-olds at identifying the function of an object not being used for its original purpose (German & Johnson, 2002) and generating potential uses outside of the design parameters of an object in a functional fluency task (Defeyter, Avons, & German, 2007). These differences are understood to arise because younger children activate more information regarding the object and possess weaker selection processes—ideas that clearly relate to our proposal that early concepts develop with minimal influence from semantic control.

### Relation to research on executive control of other domains

The current research indicates that semantic control develops later than semantic representation processes (although perhaps earlier than domain-general executive control) and demonstrates why this is advantageous. This rationale could be specific to semantic control and not the executive control of other domains. However, the ideas motivating our approach are closely connected to a recent proposal about the computational trade-offs that motivate the need for control in the first place (Musslick & Cohen, 2021), which in turn relates to recent insights about multi-task learning from computer science. Briefly, any system that must learn mappings from various input to various output channels can do so either via separate, independent pathways that each acquire their own representations, or via a shared substrate that learns a common representation for different mappings (or some blend of these possibilities). When different input/output mappings share common structure, there is a computational benefit to employing a common representation, as learning about one task will promote generalisation to and thus more rapid learning of other tasks. This is the “multi-task learning” insight that allows for zero- or few-shot learning in a variety of AI applications, including machine translation (e.g., Johnson et al., 2017) and game-playing systems (e.g., Hu, Lerer, Peysakhovich, & Foerster, 2020). With a shared representation, however, the system can only compute one input/output mapping at a time, requiring a control system to determine which inputs and outputs should engage the common representation in a given context. Moreover, different tasks must be performed in serial, as the effort to perform two different tasks in parallel will produce interference in the shared representation. Thus, capacity limits on multi-tasking arise precisely when different tasks share a representational substrate and control is needed to serialise the tasks and prevent interference. Conversely, if different input/output mappings share little structure, there is no learning benefit to using a common representation, and the system can as easily acquire separate, independent representations for each task. This avoids the need for seriality and control, since all mappings can be computed in parallel, but exerts a cost on learning since the representations in each pathway must be acquired separately. Thus, the canonical distinction between controlled, serial processing on one hand, and automatic, parallel processing on the other, represent different optimisations of a trade-off between cost of learning on the one hand versus cost of requiring serial control on the other.

Through this lens, semantic cognition can be viewed as the ultimate multi-task learning problem. As argued in the Introduction, the discernment of conceptual structure requires sensitivity to information distributed across multiple representational modalities, and across different tasks and learning episodes. The benefit of such learning is precisely that emphasised by Musslick et al. (2021) for controlled systems: use of a common conceptual representation promotes broad generalisation across items, tasks and situations. Thus, semantic cognition is a key domain where control and serial processing are required—explaining, for instance, why it is difficult to simultaneously read a book and listen to a podcast, or understand what someone is saying while you are also speaking. This perspective suggests that tasks in other cognitive domains requiring controlled, serial processing may do so because they adopt a shared representation, in which case the delayed maturation of executive control may likewise benefit learning in those domains. Future work should thus consider whether delayed maturation of domain-general executive function likewise benefits learning in other domains that also exploit shared representations.

## Methods

### Part 1: Simulations

We conducted simulations with the recent model of controlled semantic cognition from Jackson et al., (2021; see Figure 1), a deep variant of the hub-and-spokes framework in which the propagation of activation through the semantic representation network is constrained by inputs from a prefrontal “control” system. Items in the environment are represented as distributed patterns across three modality-specific input/output channels (e.g., vision, words, and action).

From input in any given channel, the model learns to activate patterns in other channels appropriate to the current task context. For instance, this may simulate a picture naming task, if the model receives direct input via the “visual” channel for a given concept, and must activate the corresponding pattern for that concept in the “words” layer, without activating the associated pattern in the “action” layer. Alternatively, it could simulate production of an associated gesture given an object’s name, if the model receives input in the “word” layer and must generate the associated pattern across “action” units, without activating “vision” units, and so on. Tasks are represented with units that locally indicate which modalities are involved—for instance, in a task taking input in M1 and producing output in M2, both the M1 and M2 units in the control layer are active. These then send connections directly to units across the deep hub-and-spokes network. To understand how control influences the development of semantic representations in this setup, we manipulated (1) the timing and speed with which controlled semantic behaviour arises and (2) the connectivity between control units and the rest of the semantic network.

#### Construction of model environment

The model environment is the same as in Jackson et al., (2021), designed to assess how well each model layer can encode cross-modal similarity structure through conceptual abstraction across sensory modalities and contexts. It includes 16 concepts, each possessing a unique activation pattern across 12 feature units in each of the 3 modality-specific information channels (see Figure 1.B.). While this environment is relatively simple, it allows us to be explicit about the nature of the structure within and across modality-specific spoke regions. The patterns were designed so that (1) the pairwise similarity structure encoded in a given modality was uncorrelated with the similarities encoded in each other modality, but with (2) a small number of features expressing common structure *across* the three modalities. This scheme captures the common insight that perceptual similarities in a given modality often differ both from the “surface” similarities in other modalities and from “deep” conceptual similarity structure—for instance, a pear and a lightbulb have similar shapes but are conceptually distinct and also have very different verbal descriptors and associated action plans. The inclusion of features that express the same structure across modalities ensures that the full multimodal structure differs from that apparent in any single channel, and captures the common intuition that conceptual structure emerges from patterns of covariation across stimulus modalities and task contexts. Note that while the unimodal and multimodal structures are orthogonal, the presence of both unimodal and multimodal structures within each modality means that the multimodal conceptual structure is not orthogonal to the information available in each sensory domain: there are features in each modality that are predictive of features in other modalities (those encoding the multimodal structure), as well as some that are only predictive of features within this modality (those encoding a unimodal structure). Thus, if the learned patterns of activation arising across a given layer express inter-item similarities highly correlated with the full multimodal structure and independent of the current task context, that layer can be understood as encoding “deep” conceptual structure.

#### Architectural details

The architecture of the semantic network model (Figure 1) was shown in prior work to best promote both learning of the latent context-independent structure present across modalities and context-sensitive responding. We describe the key parameters briefly here; for the full rationale see Jackson et al. (2021). Twelve feature units in each modality (the “Spokes” layers) employed a sigmoid activation function and possessed a fixed bias of -3, thus adopting low activations in the absence of input. Each feature unit also received a direct positive connection from a corresponding input unit, with a fixed value of +6. Direct perception of an item’s features was simulated by turning on the associated input units which, in combination with the negative biases, provided a total net input of +3 to the corresponding feature units.

These feature units were reciprocally connected to two layers of hidden units, as shown in Figure 1. Each spoke was fully and reciprocally connected to a distinct set of 14 hidden units in the Intermediate Layer, and each such layer was reciprocally connected with a common Hub Layer (including 18 units) using a sparsity of approximately 0.6349. This sparsity level was used for consistency with the prior paper, where it was necessary to match the total number of connections across different network architectures. Thus, the Hub Layer is a multimodal hub mediating between all sensory modalities, whereas regions of the Intermediate Layer support relatively modality-specific processing. Additionally, a small number of ‘shortcut connections’ project directly from the spokes to the hub, bypassing the Intermediate Layer, analogous to sparse long-range connectivity observed in the temporal lobe. Each region also has self-connections at full sparsity in the hub and spoke layers and half sparsity within each Intermediate Layer region.

When provided with an item’s partial input in one modality, the “mature” (i.e., fully trained) model must activate, across the Spoke Layer, only properties that are both true of the item and relevant to the current task context. Input and output patterns consist of the features of the relevant concept within a single modality (M1, M2, M3), resulting in 9 different ‘tasks’ as each of the three modalities may be the input or output modality (including mappings from one modality to itself, for instance in a task where the participant views a visual image and must “hold it in mind” after the stimulus vanishes, or where a word is repeated). All other input units are set to 0 and all other output units have targets of 0. The input and output modalities relevant to a given trial are specified by the context signal coded in a ‘Control Layer’ possessing 3 units, each corresponding to a single modality. A control unit is active when the associated modality is relevant to the current task as either input, output, or both. Trainable unidirectional weights from the control layer to the rest of the network thus allow the model to gate the flow of activity on the basis of the context signal so that only task-relevant properties become active in the spokes. In simulation 1, the control connections project to every layer of the semantic network allowing exploration of the relative time-course of semantic control and representation with fewer assumptions of the precise connectivity. However, metabolic and packing constraints in the brain make long-range white matter connections costly and as such this full connectivity is not biologically plausible. As such we can assume connectivity is more restricted and compare the effect of the control connections projecting selectively to the Spokes Layer, the first layer of hidden units, or the hub (simulation 2). In the prior work, the connection to the hub severely limited conceptual abstraction ability, suggesting a connection to more peripheral regions is more likely. However, these findings may differ if control is introduced later, therefore both factors are assessed in parallel here. In simulation 2, these connections were applied probabilistically, to equate the number of connections from the Control Layer to the rest of the model (matching the 18 connections possible in the Control to Hub Layer version).

#### Maturation of cognitive control

Within this framework we aimed to assess whether delayed maturation of control promotes better learning of conceptual (i.e., full context-insensitive, crossmodal) similarity structure and/or shorter time to mature performance. To test this, the model was trained for an initial *control-free* period during which the control units always adopted values of zero and the model was trained with targets on *all* of an item’s associated properties across all modalities. This is the typical framework for conceptual models without control requirements; the models are trained to produce the full set of output features, receiving feedback on each feature. The same training was provided to the initial models assessing multimodal conceptual abstraction without context-sensitive behaviour in Jackson et al., (2021). *Controlled* training items were then introduced into the environment, with gradually increasing frequency, where control units indicating the current task were activated for each learning pattern and the model was trained to activate only those units true of the item and relevant to the task (e.g., only visual and word units for picture naming).

To understand how the onset of control impacted learning, we varied (a) the *maturational delay*, that is, the duration of the training period without control, ranging from 0 to 5000 epochs in 1000 epoch increments, (b) the *transition function* describing how the frequency of controlled learning trials increased over time (linear or sigmoidal), and (c) the size of the *transition window* over which controlled trials ramped up from 0-100% of training trials (500, 1000, or 2000 epochs). The maturational delay indicates the number of epochs from the onset of training to the midpoint of the transition window (where there is an equal probability of selecting an item with and without control demands). Varying the transition window changed the number of epochs before the probability of sampling a controlled item shifted, but did not change the total number of shifts (which was always 10). We also trained a comparison model in which control was never introduced, as a reference point for interpretation of the other models. Similarly, a model with the instant addition of full control requirements was included for comparison only (see Supplementary Materials 1). Note that that there are 48 total training patterns in the control-free period (inputs for 16 items in each of 3 modalities) whereas there are 144 total training patterns in the controlled period (9 different tasks for 16 items). Each epoch included 48 training patterns sampled stochastically from amongst all patterns (with and without control) according to their specified probabilities. This resulted in the same number of learning experiences (48) and weight updates (one) per epoch with or without control, allowing for a direct comparison of training time. Control trials were introduced by gradually changing the probability of sampling a control versus no control pattern from 0 to 1 according to either the linear or sigmoidal ramping function implemented across a transition window of 500, 1000, or 2000 epochs.

#### Simulation details

All simulations were conducted within the Light Efficient Network Simulator (LENS, version 2.63) software (Rohde, 1999) with identical parameters to Jackson et al., (2021), except that we sampled initial weights from a narrower uniform range (−0.2 to 0.2) to promote a higher level of consistency across runs. The models were fully recurrent continuous-time networks using a temporal integration constant of 0.25, with activity unfolding across 24 time-steps, and input provided in the first 12. Target values were applied throughout. Training used stochastic gradient descent to minimise cross-entropy error loss using a learning rate of .001, weight decay of .0001, and no momentum. All models were trained to the same performance criterion: the model must generate the correct, task- and item-specific pattern of activation across all output units to within 0.2 of their target values. This means that properties true of an item but irrelevant to the task must be correctly inhibited; properties appropriate to the task but not true of the item must likewise be inhibited; and properties both relevant to the task and true of the item must be activated, meeting the central challenges of controlled semantic cognition. Training halted when all output units met this criterion.

#### Assessment metrics

To understand how the connectivity and maturation of control impact simulated conceptual development, we considered two metrics. The first is *training time,* measured as the number of learning epochs required before the model generates the correct task-specific outputs for all *controlled* patterns in the environment. The second is the *conceptual abstraction score*, defined as the degree to which learned representations in the trained model capture the full similarity structure of the environment abstracted across items, learning episodes, contexts and modalities (Jackson et al., 2021; McRae et al., 2005; Rogers & McClelland, 2004). To measure conceptual abstraction, we first computed a *target conceptual similarity matrix* containing the pairwise correlations between all items in the model environment based on their full set of properties across modalities. For each hidden layer in a given model we also computed a *representational similarity matrix* containing pairwise correlations for all items in the evoked pattern of activation generated across units in the layer by each item input (at the final time point within a trial). Conceptual abstraction was defined as the Pearson correlation between the representational similarity matrix and the target matrix. We report this for the Hub Layer only since it always had the highest score, but values for the Intermediate Layer are reported in Supplementary Materials 2.

For each simulation condition we trained 80 models differing only in their weight initialisation to calculate the conceptual abstraction scores and 25 models to compute training time. The same sample sizes were used (here and throughout) as in Jackson et al., (2021) which are far higher than typical modelling investigations as large effect sizes are expected in simulation data. To understand whether and how the delayed onset of control impacted these metrics, we first used an ANOVA to assess how they were affected by three factors governing the transition to fully controlled trials—maturational delay (i.e., the duration of the period without control), the transition function governing the ramp-up of control, and the length of the transition window.

We then assessed the nature of the effect of maturational delay on both metrics using independent-samples t-tests to contrast each delay period to the baseline condition in which the model was trained only with control trials from the onset of training (as in Jackson et al., 2021). Where necessary (determined using Levene’s test) t-tests were corrected for unequal variance between the groups, and all p-values were Bonferroni-adjusted for 5 comparisons. For interpretation of the size of these effects, Cohen’s d effect sizes and 95% confidence intervals are reported. For ANOVAs partial eta squared is reported. Code to run all simulations and process the results is available at github.com/JacksonBecky/developmental.

### Part 2: Meta-analysis of Developmental Data

We conducted a literature review seeking match-to-sample studies employing either thematic or superordinate taxonomic matching or both. A match-to-sample task is defined as presentation of a single item as a probe or standard for comparison, and two or more targets requiring an active choice to decide which is related to the standard in some way (indicated via pointing, picking up an object, or a spoken response; preferential-looking paradigms were excluded). Stimuli could be pictures, words or objects. The measure of interest was the frequency of choosing the taxonomic or the thematic match; studies reporting reaction time or individual differences only were excluded. We then considered two kinds of studies:

1. *Low control*, where the target matched the sample either taxonomically or thematically, and remaining options were semantically unrelated to the sample. These trials thus indicated whether the participant possessed taxonomic or thematic knowledge sufficient to choose the target without having to select what kind of semantic information was relevant for the task.
2. *High control,* where all trials have both a thematic and taxonomic match but the instructional context indicates which kind of information should drive the selection. These trials thus require both the relevant semantic knowledge and the ability to select the correct kind of information for the task.

Studies were initially identified from two sources: the list of developmental studies of taxonomic and thematic structures in Mirman et al., (2017) and a Web of Knowledge search with any two of the terms ‘taxonomic’, ‘thematic’ and ‘perceptual’ or with the term ‘noun-category bias’, performed in January 2021 and including full papers from any year. This resulted in an initial screening of 383 studies. The references of all included studies were also screened for inclusion. Studies were excluded if participants were explicitly rewarded for taxonomic vs. thematic choices, or if the studies intentionally weighted the taxonomic and thematic responses to be stronger or weaker than each other. Studies could include additional response options which were factored out to assess only the contrast of interest. Studies with more than two choices were corrected to a chance level of 0.5. All studies assessed healthy participants without neurostimulation during childhood or young adulthood (studies with mean ages over 55 were excluded to avoid possible effects of ageing). High control studies were sorted based on their instructions as described above. Studies using ambiguous instructions unlikely to strongly weight the context toward taxonomic vs. thematic cues in adults (e.g., choose ‘the most related’ or ‘more similar’ item) were excluded. For the low control studies we did not further sort based on instructions as any semantic relationship cue should lead participants to choose the correct (taxonomic or thematic) target over an unrelated concept.

For each study the average proportion of taxonomic or thematic responses was recorded, as well as the standard deviation (where present) and the number of participants. Studies were chunked on the basis of age (i.e., studies with an average age between 12 and 24 months were grouped as one-year-olds, etc.) from ages one to seven and adults (ages 18-55). For low control conditions we found 18 taxonomic-match experiments consisting of 639 participants across 28 data points and 17 thematic-match experiments with 657 participants across 29 data points. For high control conditions, we found 20 experiments employing a clear thematic cue, consisting of 805 participants over 35 data points, and 22 experiments employing a clear taxonomic cue, consisting of 561 participants over 29 data points. Thus, data from 2662 participants contributed to the assessment of semantic control and representation ability across development. See Supplementary Materials 3 for full details of the studies included at each age, including the data. The methods used throughout comply with APA ethical standards, as all data was secondary (or simulated).

Statistics were performed in SPSS (IBM, 2017) on the average proportions of taxonomic or thematic responses in each study. To determine whether each age group had evidence of a conceptual representation structure, the proportion of correct responses in the taxonomic-only and thematic-only conditions were compared to chance (0.5), using one-tailed one-sampled t-tests. To test for evidence of significant context effects, the proportion of taxonomic (vs. thematic) responding in studies with a taxonomic or thematic instruction was compared using one-tailed between-samples t-tests (as the specific expectation was that a taxonomic cue should have greater taxonomic responding). Where possible, effect sizes (g) were calculated using Hedges correction for unequal sample sizes. To fully understand the distribution of responses and for graphical purposes, the response distribution across all studies was estimated based on the reported mean and standard deviation per study. The majority of studies reported the variance. In some cases this was reported separately for different conditions and then combined. Where variance was not reported, the maximum possible standard deviation was computed. This combination of reported and maximum standard deviations provides the best estimate of the real variance whilst being conservative. However, the same pattern of results is obtained if the maximum possible standard deviation is used for all studies. These analyses were performed using R Statistical Software (v3.5.2; R Core Team, 2021) using custom built code available at github.com/JacksonBecky/developmental. For each study, the results were simulated 1000 times by drawing a number of samples equal to the number of participants in the study from a normal distribution with the reported mean and standard deviation. These simulated data points were used to compute a grand mean (across all studies and simulation iterations, which is weighted by the number of participants), which is displayed in the plot. The .025 and .975 quantiles of this distribution were used to compute the 95% confidence intervals of the mean across studies.

Where the confidence intervals for thematic or taxonomic over unrelated selections reliably exclude 0.5, this further supports the conclusion that this form of knowledge is present above chance level. Where the confidence intervals for the taxonomic and thematic cues reliably exclude each other, this further supports the presence of context-sensitive responding to differential cues.

#### Transparency & openness

We report how we determined our sample size, all data exclusions, all manipulations and all measures in all studies within this article. We follow JARS (Kazak, 2018). All data are available in the Supplementary Materials or can be generated with the code provided. All code is available at github.com/JacksonBecky/developmental. Data were generated using LENS (v2.63; Rohde, 1999) and analysed using Matlab (The MathWorks, 2018), SPSS (v.25; IBM, 2017) and R Statistical Software (v3.5.2; R Core Team, 2021). The study design and analyses were not preregistered.

## Acknowledgements

This work was supported by a British Academy Postdoctoral Fellowship awarded to R.L.J. (no. pf170068), a programme grant to M.A.L.R. from the Medical Research Council (grant no. MR/R023883/1), an Advanced Grant from the European Research Council to M.A.L.R. (GAP: 670428) and Medical Research Council intramural funding (no. MC_UU_00005/18).

## Data & Code Availability

All data are available in the Supplementary Materials or can be generated with the code provided. All code is available at github.com/JacksonBecky/developmental.

For the purpose of open access, the author has applied a Creative Commons Attribution (CC BY) license to any Author Accepted Manuscript version arising from this submission.

