## Supplementary Materials for "Late maturation of semantic control promotes conceptual development"

Supplementary Note 1: The Effects of a Maturational Delay in Control when Instantly Adding Control

The length of a developmental period without control was manipulated with the instant addition of control in the model with full connectivity from control (see Supplementary Figure 1). Although not a viable hypothesis as to how control is added within the brain, this provides a comparison for other control protocols. The pattern of effects was the same as the gradual addition protocols. Conceptual abstraction was found to vary based on the length of the maturational delay ($F(5,474)=17.301$, $p<.001$), with the difference from the baseline only reaching significance at the final time point (control throughout *vs.* 1000 epochs; $t(158)=-0.824$, $p=1$, CI=-.020, .008; *vs.* 2000 epochs; $t(158)=-1.725$, $p=.433$, CI=-.026, .002; *vs.* 3000 epochs; $t(158)=0.256$, $p=1$, CI=-.013, .017; *vs.* 4000 epochs; $t(158)=1.685$, $p=.470$, CI=-.002, .027; *vs.* 5000 epochs; $t(158)=6.036$, $p<.001$, CI=.032, .064). Effect sizes were small until this time (control throughout *vs.* 1000 epochs; d=.130; *vs.* 2000 epochs; d=.273; *vs.* 3000 epochs; d=.041; *vs.* 4000 epochs; d=.266; *vs.* 5000 epochs; d=.950). Training time also differed by the length of the developmental period ($F(5,144)=26.639$, $p<.001$). A developmental period without control resulted in much faster learning, with longer delays having larger effects (control throughout *vs.* 1000 epochs; $t(48)=2.146$, $p=.185$, CI=51.95, 1597.25, d=.607; *vs.* 2000 epochs; $t(48)=3.226$, $p=.011$, CI=451.41, 1944.67, d=.913; *vs.* 3000 epochs; $t(48)=5.860$, $p<.001$, CI=1348.20, 2756.52, d=1.658; *vs.* 4000 epochs; $t(44.973)=9.074$, $p<.001$, CI=2341.64, 3677.72, d=2.567; *vs.* 5000 epochs; $t(40.791)=9.705$, $p<.001$, CI=2400.25, 3661.91, d=2.745).

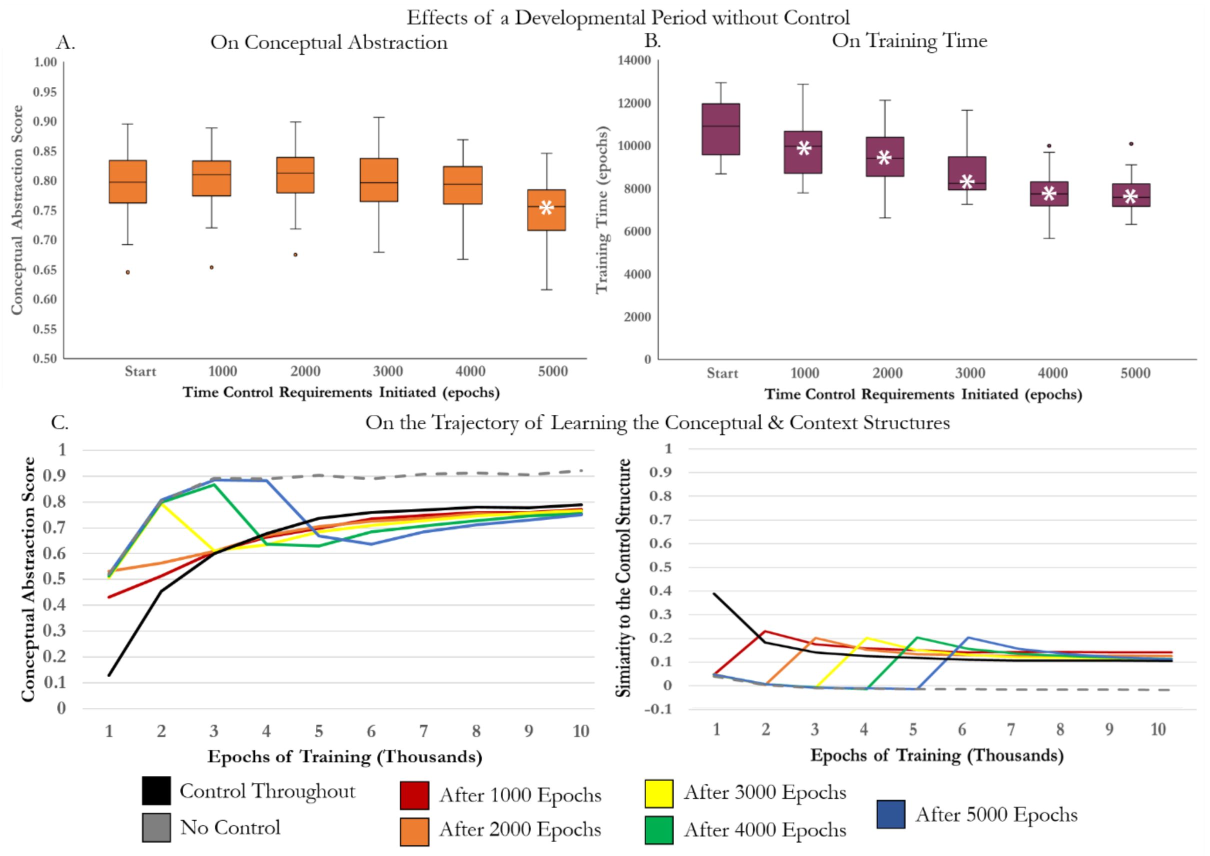


*Supplementary Figure 1. The effects of a developmental period without control on the model with full connectivity from control. Control is added in full from the start, or after 1000, 2000, 3000, 4000 or 5000 epochs. A. The effects of a developmental period without control on the conceptual abstraction score. B. The effect of a developmental period without control on the time taken to train the model. Bars signify the median value and the 25$^{th}$ and 75$^{th}$ percentile values. White asterisks signify a significant effect of the delay in adding control in contrast to the inclusion of control requirements from the start of training. C. The effects of a developmental period without control on the evolution of the conceptual abstraction score and the representation of the context signal during training. The mean conceptual abstraction score (left) and the similarity to the control structure (right) are displayed after every 1000 epochs of training. Both are based on the representations in the Hub Layer alone. The trajectory of conceptual abstraction when control is never added is shown with a dashed line as this model would never meet the requirement of context-sensitive behaviour that is central to the human semantic system.*

### Supplementary Note 2: The Effects of a Maturational Delay in Control on Intermediate Layer Representations

As the deep hub region is critical for representing the context-independent representation structure (denoting conceptual abstraction), the focus in the main text is the Hub Layer representations. However, we can also assess the similarity to this structure within the Intermediate Layer, collapsing across the six different protocols for gradually adding control. This displays a decrease with longer developmental periods without control ($F(5, 8634)=441.694$, $p<.001$; control after 1000 epochs; $t(2878)=12.730$, $p<.001$, $d=.474$, CI=.023, .031; control after 2000 epochs; $t(2874.404)=24.751$, $p<.001$, $d=.922$, CI=.047, .055; control after 3000 epochs; $t(2868.605)=31.547$, $p<.001$, $d=1.176$, CI=.061, .069; control after 4000 epochs; $t(2867.348)=35.744$, $p<.001$, $d=1.332$, CI=.069, .077; control after 5000 epochs; $t(2874.927)=38.016$, $p<.001$, $d=1.417$, CI=.075, .083; see Supplementary Figure 2). Displaying the change in the similarity to the conceptual and contextual similarity matrices over learning (in the logistic protocol for adding control, applied over 1000 epochs) helps interpretation of the changes in the deeper hidden layer. The Intermediate Layer learns representations that are more sensitive to context than the hub. If control requirements are present throughout training both hidden layers initially learn the context structure. However, if control is initiated after a delay, both structures initially learn some conceptual structure, allowing the hub to learn conceptual information faster. When control is instantiated, the Intermediate Layer quickly becomes more context-sensitive and this effect persists throughout time (unlike the hub which can recover conceptual information).

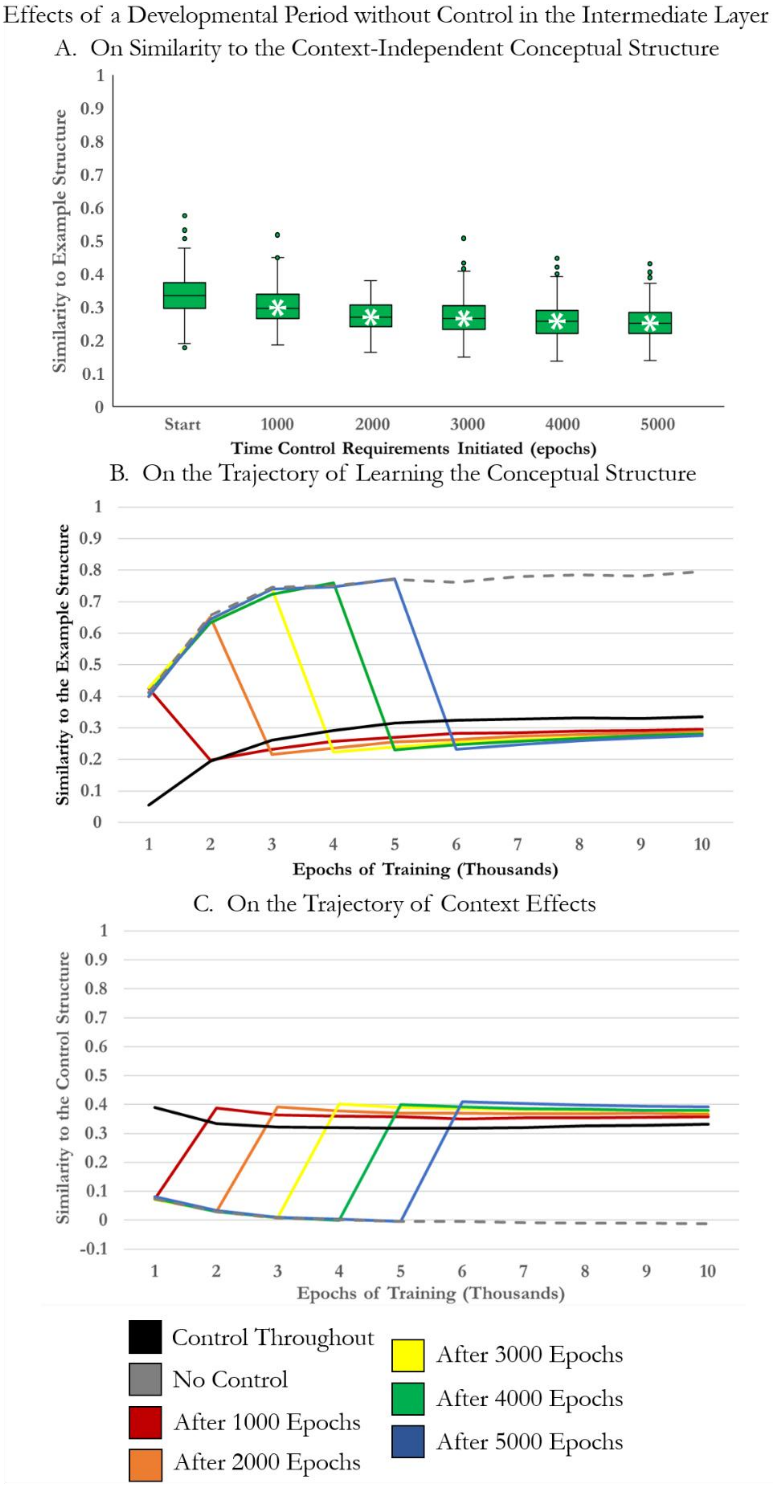

*Supplementary Figure 1. The effects of a developmental period without control on Intermediate Layer representations. Control is present from the start, or added gradually around 1000, 2000, 3000, 4000 or 5000 epochs. A. The effects on the similarity of the Intermediate Layer representations to the conceptual similarity matrix (collapsed across all protocols for gradually adding control). Bars signify the median and the 25th and 75th percentile values. White asterisks signify a significant effect of the maturational delay in contrast to requiring control from the start of training. B&C. The effects of a developmental period without control on the evolving similarity to the conceptual and contextual similarity matrices in the Intermediate Layer after every 1000 epochs of training. Control is added via the logistic protocol implemented across 1000 epochs. The trajectory of learning when control is never added, is shown with a dashed line as this model would never meet the requirement of context-sensitive behaviour that is central to the human semantic system. As the Intermediate Layer is more context-sensitive than the hub, it always has high similarity to the contextual and low to the conceptual matrix once control is added. However, in a developmental period it temporarily assumes a role in context-insensitive conceptual representation, which allows greater conceptual information to be represented in the deep multimodal hub at an earlier stage of learning.*

### Supplementary Note 3: Behavioural Data Details

#### Taxonomic vs. thematic studies with taxonomic instructions

| Paper | Year | Study | Age (years) | Proportion (taxonomic) | n |
|---|---|---|---|---|---|
| Imai et al. | 1994 | Exp 1 | 3 | 0.32 | 15 |
| | | | 5 | 0.64 | 15 |
| | | | Adult | 0.97 | 15 |
| Markman & Hutchinson | 1984 | Exp 2 | 4 | 0.687 | 15 |
| Markman & Hutchinson | 1984 | Exp 3 | 4 | 0.645 | 20 |
| Imai et al. | 2010 | Exp 2 | 3 | 0.71 | 33 |
| | | | 5 | 0.732 | 32 |
| Waxman, Philippe, et al. | 1999 | Exp 1 | 3 | 0.908 | 12 |
| Waxman, Philippe, et al. | 1999 | Exp 2 | 3 | 0.525 | 9 |
| Golinkoff, Shuffbailey, et al. | 1995 | Exp 1&5 | 3 | 0.658 | 16 |
| | | | 7 | 0.952 | 14 |
| | | | Adult | 1 | 9 |
| Golinkoff, Shuffbailey, et al. | 1995 | Exp 2 | 3 | 0.524 | 12 |
| Golinkoff, Shuffbailey, et al. | 1995 | Exp 3 | 4 | 0.5 | 12 |
| Poulin-Dubois, Frank, et al. | 1999 | Exp 1 | 2 | 0.5346 | 31 |
| Poulin-Dubois, Frank, et al. | 1999 | Exp 2 | 2 | 0.4449 | 48 |
| Waxman, Senghas, et al. | 1997 | Exp 1 | 3 | 0.66 | 30 |
| Waxman, Senghas, et al. | 1997 | Exp 2 | 4 | 0.87 | 15 |
| Waxman, Senghas, et al. | 1997 | Exp 3A | 6 | 0.7 | 15 |
| Waxman, Senghas, et al. | 1997 | Exp 3B | 6 | 0.67 | 10 |
| Waxman & Kosowski | 1990 | Exp 1&3 | 2 | 0.72 | 15 |
| | | | 3 | 0.69 | 15 |
| Baldwin et al. | 1992 | Exp 2 | 4 | 0.49 | 10 |
| Liu et al. | 2001 | Exp 2 | 5 | 0.63 | 22 |
| Liu et al. | 2001 | Exp 3 | 4 | 0.85 | 16 |
| Liu et al. | 2001 | Exp 6 | 3 | 0.53 | 16 |
| | | | 4 | 0.92 | 18 |
| Tare & Gelman | 2010 | Exp 1 | Adult | 0.99 | 35 |
| Tare & Gelman | 2010 | Exp 2 | 4 | 0.77 | 36 |

Taxonomic vs. thematic studies with thematic instructions

| Paper | Year | Study | Age (years) | Proportion (taxonomic) | n |
|---|---|---|---|---|---|
| Greenfield & Scott | 1986 | Exp 1 | 3 | 0.33 | 12 |
| | | | 4 | 0.15 | 12 |
| | | | 5 | 0.07 | 12 |
| | | | 6 | 0.08 | 12 |
| | | | 7 | 0.26 | 12 |
| Blaye & Bonthoux | 2001 | Exp 1 | 5 | 0.52 | 69 |
| Imai et al. | 1994 | Exp 1 | 3 | 0.29 | 15 |
| | | | 5 | 0.3 | 15 |
| | | | Adult | 0.36 | 15 |
| Waxman & Namy | 1997 | Exp 1&3 (goes with) | 2 | 0.56 | 16 |
| | | | 3 | 0.54 | 16 |
| | | | 4 | 0.44 | 16 |
| Waxman & Namy | 1997 | Exp 1&3 (goes best) | 2 | 0.51 | 16 |
| | | | 3 | 0.46 | 16 |
| | | | 4 | 0.21 | 16 |
| Waxman & Namy | 1997 | Exp 2 (goes with) | 3 | 0.55 | 8 |
| | | | 4 | 0.46 | 8 |
| Waxman & Namy | 1997 | Exp 2 (goes best) | 3 | 0.52 | 8 |
| | | | 4 | 0.19 | 8 |
| Lin & Murphy | 2001 | Exp 3 | Adult | 0.27 | 18 |
| Lin & Murphy | 2001 | Exp 5 | Adult | 0.3 | 20 |
| Osborne & Calhoun | 1998 | Exp 1 | 4 | 0.614 | 41 |
| Osborne & Calhoun | 1998 | Exp 2 | 4 | 0.675 | 41 |
| Osborne & Calhoun | 1998 | Exp 3 | 4 | 0.578 | 41 |
| Luariello et al. | 1992 | Exp 3 | 4 | 0.329 | 20 |
| | | | 7 | 0.229 | 20 |
| Saalbach & Imai | 2007 | Exp 1 | Adult | 0.353 | 47 |
| Skwarchuk & Clark | 1996 | Exp 1 | Adult | 0.2389 | 24 |
| Skwarchuk & Clark | 1996 | Exp 2 | Adult | 0.2653 | 24 |
| Skwarchuk & Clark | 1996 | Exp 3 | Adult | 0.1819 | 24 |
| Unsworth et al. | 2005 | Exp 2 | Adult | 0.503 | 67 |
| Ware et al. | 2017 | Exp 1 | 5 | 0.39 | 23 |
| | | | 6 | 0.19 | 22 |
| | | | 7 | 0.14 | 23 |
| Wyatt & Rabinowitz | 2010 | Exp 1 | Adult | 0.371 | 48 |

Taxonomic vs. baseline studies

| Paper | Year | Study | Age (years) | Proportion (taxonomic) | n |
|---|---|---|---|---|---|
| Fenson et al. | 1989 | Exp 1 | 2 | 0.66 | 35 |
| Landrigan & Mirman | 2018 | Exp 1 | Adult | 0.97 | 32 |
| Jackson et al. | 2015 | Pilot | Adult | 0.887 | 9 |
| Golinkoff, Shuffbailey, et al. | 1995 | Exp 2 (another dax) | 3 | 0.6 | 12 |
| Golinkoff, Shuffbailey, et al. | 1995 | Exp 2 (another one) | 3 | 0.72 | 12 |
| Fenson et al. | 1988 | Exp 1 | 2 | 0.65 | 25 |
| Scott et al | 1985 | Exp 1 | 3 | 0.75 | 24 |
| | | | 4 | 0.725 | 24 |
| | | | 5 | 0.779 | 24 |
| | | | 6 | 0.796 | 24 |
| Scott et al. | 1982 | Exp 1 | 2 | 0.83 | 16 |
| | | | 3 | 0.86 | 16 |
| | | | 4 | 0.88 | 16 |
| | | | 5 | 0.95 | 16 |
| Deahler et al. | 1979 | Exp 3 | 1 | 0.667 | 16 |
| | | | 2 | 0.771 | 32 |
| Liu et al. | 2001 | Exp 3 | 4 | 0.83 | 16 |
| Liu et al. | 2001 | Exp 6 (another dax) | 3 | 0.56 | 16 |
| | | | 4 | 0.96 | 18 |
| Liu et al. | 2001 | Exp 6 (another one) | 4 | 1 | 18 |
| Saalbach & Imai | 2007 | Exp 1 | Adult | 0.945 | 47 |
| Scott et al. | 1980 | Exp 2 (Level 3) | 4 | 0.841 | 72 |
| Nguyen & Murphy | 2003 | Exp 2 | 3 | 0.6 | 16 |
| | | Exp 3 | 4 | 0.66 | 16 |
| | | | 7 | 0.89 | 16 |
| | | | Adult | 0.89 | 16 |
| Osborne & Calhoun | 1998 | Exp 2 | 4 | 0.962 | 41 |
| Sachs et al. | 2008 | Exp 1 | Adult | 0.95 | 14 |

Thematic vs. baseline studies

| Paper | Year | Study | Age (years) | Proportion (thematic) | n |
|---|---|---|---|---|---|
| Blanchet et al. | 2001 | Exp 1 | 2 | 0.558 | 21 |
| | | | 3 | 0.682 | 20 |
| | | | 4 | 0.955 | 20 |
| Deahler et al. | 1979 | Exp 3 | 1 | 0.535 | 16 |
| | | | 2 | 0.656 | 32 |
| Fenson et al. | 1989 | Exp 1 | 2 | 0.71 | 35 |
| Golinkoff, Shuffbailey, et al. | 1995 | Exp 2 (another dax) | 3 | 0.576 | 12 |
| Golinkoff, Shuffbailey, et al. | 1995 | Exp 2 (another one) | 3 | 0.895 | 12 |
| Jackson et al. | 2015 | Pilot | Adult | 0.903 | 9 |
| Landrigan & Mirman | 2018 | Exp 1 | Adult | 0.97 | 32 |
| Liu et al. | 2001 | Exp 3 | 4 | 0.46 | 16 |
| Liu et al. | 2001 | Exp 6 (another dax) | 3 | 0.53 | 16 |
| | | | 4 | 0.67 | 18 |
| Nguyen & Murphy | 2003 | Exp 2 | 3 | 0.6 | 16 |
| Nguyen & Murphy | 2003 | Exp 3 | 4 | 0.6 | 16 |
| | | | 7 | 0.83 | 16 |
| | | | Adult | 0.89 | 16 |
| Osborne & Calhoun | 1998 | Exp 2 | 4 | 0.924 | 41 |
| Saalbach & Imai | 2007 | Exp 1 | Adult | 0.966 | 47 |
| Sachs et al. | 2008 | Exp 1 | Adult | 0.97 | 14 |
| Scott et al. | 1980 | Exp 2 (Level 3) | 4 | 0.903 | 72 |
| Scott et al. | 1982 | Exp 1 | 2 | 0.88 | 16 |
| | | | 3 | 0.94 | 16 |
| | | | 4 | 0.96 | 16 |
| | | | 5 | 1 | 16 |
| Scott et al | 1985 | Exp 1 | 3 | 0.762 | 24 |
| | | | 4 | 0.767 | 24 |
| | | | 5 | 0.792 | 24 |
| | | | 6 | 0.792 | 24 |
